# A comparative analysis of the ontogeny of syngnathids (pipefishes & seahorses) reveals how heterochrony contributed to their diversification

**DOI:** 10.1101/2022.08.08.503137

**Authors:** Ralf Friedrich Schneider, Joost Marten Woltering, Dominique Adriaens, Olivia Roth

## Abstract

**Background:** Syngnathids are a highly derived and diverse fish clade comprising the pipefishes, pipe-horses and seahorses. They are characterized by a plethora of iconic traits that increasingly capture the attention of biologists, including geneticists, ecologists and developmental biologists. The current understanding of the origins of their derived body plan is, however, hampered by incomplete and limited descriptions of the early syngnathid ontogeny.

**Results:** We provide a comprehensive description of the development of *Nerophis ophidion*, *Syngnathus typhle*, *Hippocampus erectus* from early cleavage stages to release from the male brooding organ and beyond, including juvenile development. We comparatively describe skeletogenesis with a particular focus on dermal bony plates, the snout-like jaw morphology, and appendages.

**Conclusions:** This most comprehensive and detailed account of syngnathid development to date suggests that convergent phenotypes (e.g. reduction and loss of the caudal fins), likely arose by distinct ontogenetic means in pipefish and seahorses. Comparison of the ontogenetic trajectories of *S. typhle* and *H. erectus* provides indications that characteristic features of the seahorse body plan result from developmental truncation. Altogether this work provides a valuable resource and framework for future research to understand the evolution of the outlandish syngnathid morphology from a developmental perspective.

**Bullet points:**

- Syngnathids (pipefish, pipe-horses and seahorses) are a highly derived and diverse fish taxon and many of their most iconic traits – and these traits’ variation across species - develop during their early ontogeny.
- This study provides a detailed description and staging table of the entire pre-release development (and beyond) of three distinct syngnathids (two phylogenetically distinct pipefishes and a seahorse) with a focus on the skeletal elements forming the cranium, axial skeleton, appendages and the plate armor.
- The data provided in this study suggests that developmental heterochrony, i.e. alterations in the rate and timing of developmental processes between lineages, might be a central evolutionary process shaping the morphological diversity found in syngnathids today.

## Introduction

Syngnathids (pipefishes, pipe-horses and seahorses) are a globally distributed taxon of highly derived fishes that evolved a plethora of bizarre and unique phenotypes. Examples are the elongated, fin-reduced bodies of the pipefishes,^1^ the leaf-mimicking appendages of the leafy seadragons,^2,3^ the curved body posture that permits upright swimming in seahorses ^4^, and of course male pregnancy - a unique trait shared amongst syngnathids.^5,6^ These and other outlandish novelties of the syngnathid lineage attract an ever-increasing scientific attention.^7,8^ For instance, recent studies addressed how the syngnathids’ tube-like jaws and angled head posture facilitate pivot feeding,^9,10^ or investigate the mechanical basis for the flexible and strong prehensile tail.^11,12^ A full comprehension of the evolutionary history of lineage specific morphological novelties, that is beyond their adaptive significance or adult physiology, requires knowledge of how these traits form during embryonal development. In case of the syngnathids, several factors complicate the study of their ontogeny. Firstly, laboratory work on syngnathids requires a dedicated marine facility and breeding only occurs with optimized husbandry. Alternatively, embryonic material can be collected in the field but typically this is cumbersome with respect to culturing and documentation. Secondly, syngnathids have a reproductive strategy involving paternal care whereby the embryos are fertilized and develop inside a ventral belly-groove or pouchlike brooding organ of the father. Many syngnathids (e.g., seahorses and *Syngnathus* pipefishes) produce embryos that upon release already possess the adult body plan and thus do not pass through a specialized free-living larval stage – i.e., they qualify as direct developers.^13–15^ Therefore, the most informative developmental stages need to be collected from the brooding organ through more or less invasive procedures (depending on the species and type of brooding organ, see below). Thus, to study the developmental biology of specific morphological features, such as the elongated snout or fin loss in pipefishes, their pre-release development must be considered.

Syngnathid embryonal and juvenile development has been the subject of several studies: Novelli et al.^16,17^ provided a detailed description of the chondro- and osteoskeletal development in juvenile Short-snouted seahorses (*Hippocampus hippocampus)* throughout the first month post-release and of the inner organs in juvenile Longsnout seahorses (*Hippocampus reidi)*, confirming that loss of the spleen is an adult phenomenon. Franz-Odendaal et al.^18^ focused on post-release development of the juvenile seahorses’ head skeleton proposing that heterochronic shifts contribute to interspecific differences in head morphology. Studies investigating pre-release (i.e., embryonal) development are scarcer, and often rely upon incidental sampling with few replications, which limits the level of detail they provide (e.g., ^19–23^, but see ^24^ for approximately the second half of pre-release development, and ^25^ for *Syngnathus* kidney development). The only comparative study spanning the entire syngnathid pre-release development we are aware of describes and compares Straight-nose pipefish (*Nerophis ophidion)*, Black-striped pipefish (*Syngnathus abaster)* and Big-belly seahorse (*Hippocampus abdominalis)*.^20^ Yet only few developmental stages per species are described and developmental timing is estimated according to previous studies on related species. Furthermore, the study by Marie Azzarello^24^ focuses on the second half of pre-release development. However, due to its publication in 1990 the quality of images provided is limited to the standards of the time. Therefore, a comprehensive comparative study describing and illustrating syngnathid development remains a timely contribution to the field.

Here, we comparatively described the pre-release development of representatives of three deep syngnathid branches (Fig 1 a), the *Nerophinae* (Straight-nose pipefish, *Nerophis ophidion*), *Syngnathinae* (Broadnosed pipefish, *Syngnathus typhle*) and *Hippocampinae* (a taxon nested within the *Syngnathinae* (Fig. 1 a-d); Lined seahorse, *Hippocampus erectus*).^26,27^ Selected species feature an abundance of distinct morphological phenotypes, as they feature distinct reproductive strategies (Fig. 1 e-g). We focus on the pre-release development and add details on post-release development for some characters. Finally, we discuss the scientific value of the syngnathid model system in the context of evo-devo research.

**Figure 1:**
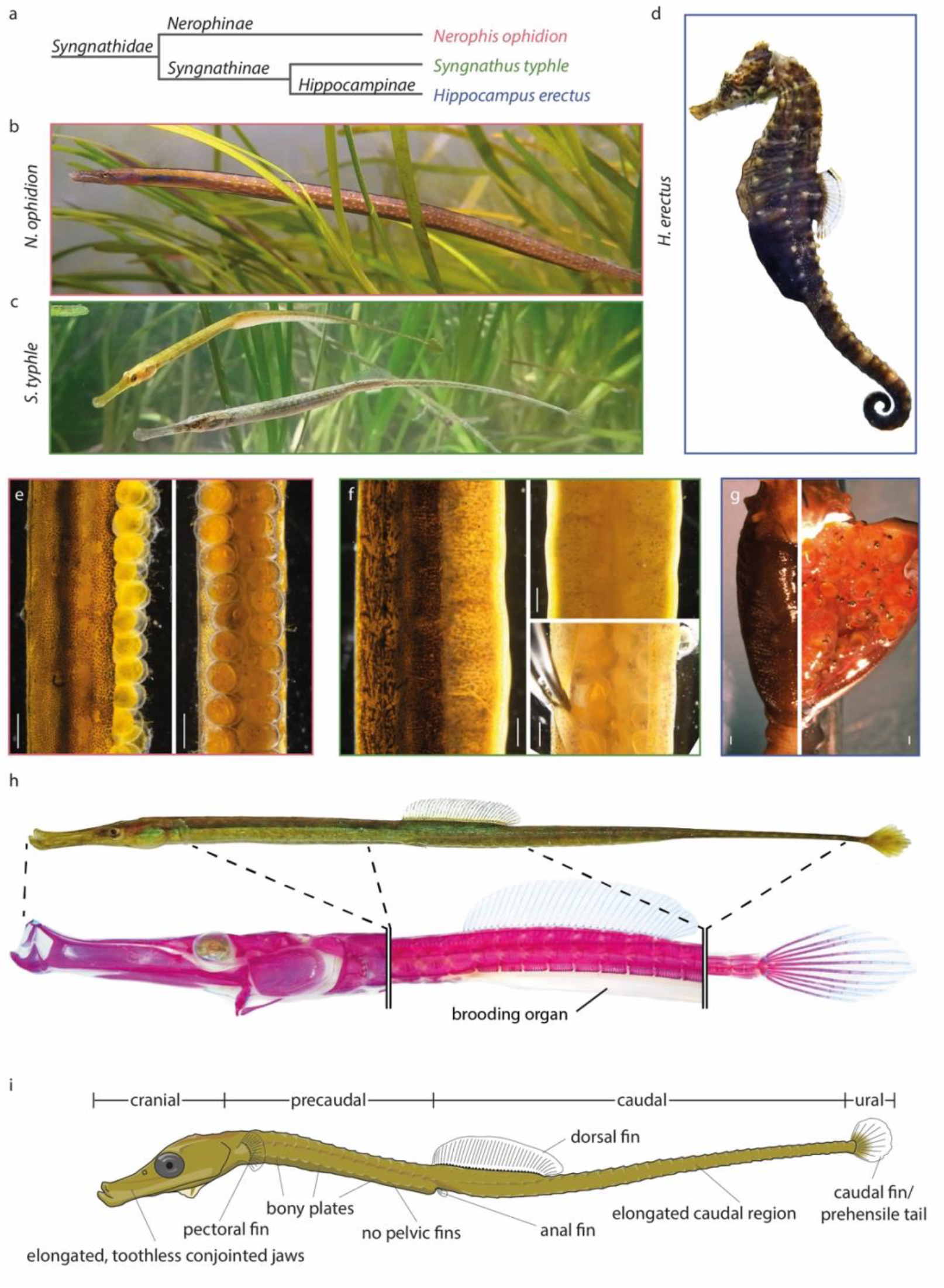
Habitus, brooding organs and body plan of examined species. (a) Cladogram of investigated species (following ^26^). (b-d) Female *N. ophidion*, Male (upper) and female (lower) *S. typhle* and male *H. erectus*, respectively. (e) Male *N. ophidion* brooding organ with eggs located along the animals’ trunk (left: lateral view; right: ventral view). (f) Male *S. typhle* brooding organs are on the animal’s tail and cover the eggs (left: lateral view; upper right: ventral view; lower right: ventral view with covering skin pulled aside). (g) Male *H. erectus* carry the developing embryos in a pouch-like brooding organ, which has a small anterior opening, on their tail (left: ventral view; right: ventral view with surgically opened brooding organ). (h) An Alizarin Red (stains ossified tissue) and Alcian Blue (stains cartilage) stain of an adult male *S. typhle* illustrates the bony armor of adult syngnathids. (i) Schematic drawing of a *S. typhle* juvenile featuring syngnathid traits focal for this study.

## Results

### Adult morphology and breeding biology: a brief overview

The three selected syngnathid species examined in this study represent three deep branches within the syngnathids radiation (Fig. 1 a), and display many of their diverse phenotypes (Fig. 1).^26^ The Straight-nose pipefish (*N. ophidion*) is a primarily European species with a preference for brackish water. It has an extremely thin and elongated body (33 precaudal and 73 caudal vertebrae; Table 1), with males typically being approximately 20cm long (with a maximum body diameter of 3-4mm) while the females are larger and can reach lengths of approximately 30cm (with a maximum body diameter of 4-6mm, not accounting for an ornamental ventral longitudinal skin protrusion; Fig. 1b; ^28^). Adult *N. ophidion* have only a dorsal fin, which contains 33 soft-rays (Table 1). Note that all syngnathids lack the fin spines present in other spiny-rayed fishes ^29^ as a result of secondarily loss. The posterior body axis does not terminate in a caudal fin but instead forms a prehensile tail used by the fish to wrap themselves around holdfasts, such as eelgrass.^30^ Females have a membrane ventrolateral along their trunk that is erected during intraspecific interactions. For syngnathids, *N. ophidion* has a relatively small head and short snout (Fig. 1).

**Table 1:** Adults’ vertebrae and ray numbers observed in studied species.

|  | *N. ophidion* | *S. typhle* | *H. erectus* |
| --- | --- | --- | --- |
| precaudal vertebrae | 33 | 19 | 12 |
| caudal vertebrae | 73 | 36 | 35 |
| ural vertebrae | 0 | 1 | 1 |
| pectoral fin rays | NA | 14 | 16 |
| dorsal fin rays | 33 | 37 | 18 |
| anal fin rays | NA | 3 | 4 |
| caudal fin rays | NA | 10 | 0 (2-5 rud.) |

The Broad-nosed pipefish (*S. typhle*) is also a European species. It has a less extremely elongated body (with 19 precaudal, 36 caudal and one ural vertebra) and males and females are more equal in size, with both rarely reaching substantially over 20cm in our sampling area of the Baltic Sea (with a maximum body diameter of approximately 1.5 cm in non-pregnant males and non-gravid females; ^28,31^; Fig. 1 c). *S. typhle* has relatively well-developed pectoral (14 rays), dorsal (37 rays), anal (3 rays) and caudal (10 rays) fins (pelvic fins are absent in all syngnathids as the result of loss of the *tbx4* gene (Table 1).^1,32^ *S. typhle* has the long tubular snout typical for syngnathids, which in this species appears laterally compressed when compared to other pipefish species.^30^ Furthermore, males have modified bony body plates with ventrolateral protrusions on the anterior portion of their tail that support the elaborate pouch-like brooding organ (Fig. 1 h).

The Lined seahorse (*H. erectus)* belongs to the subfamily of *Hippocampinae*, which have evolved an upright swimming posture, a prehensile muscular tail and the most derived form of pregnancy among syngnathids (Fig. 1 d).^33^ Males and females are similar in size; approximately 6-12 cm with uncoiled tail ^34^. However, large males typically have a deeper body due to a ventral bony trunk keel that females lack. Males also have a sealable brood pouch at the more cranial portion of their tail (Fig. 1 d). Their head (excluding snout) appears axially compressed when compared to other syngnathids (see also ^10^).

*N. ophidion*, *S. typhle* and *H. erectus* represent three main types of brooding organs that evolved in Syngnathids.^6^ *N. ophidion* has the simplest type of brooding organ, possibly representing the ancestral state for syngnathids, which runs as a ventral abdominal groove along the ventral side of the trunk (i.e. pre-caudal region). The round eggs (typically 0.8-1 mm in diameter; see also ^28^) are attached to this groove in several rows during mating (Fig. 1 e). Egg numbers are linked to male and female body size, but often exceed 100. On occasion, clutches are (partially) unfertilized, leading to a rejection of the unfertilized eggs after several days. Eggs found on wild-caught specimens are often overgrown by different types of algae and even small mussels are regularly found settling on the eggs. This does not seem to affect the survival of the eggs though (personal observation). Eggs are held in place by being partially engulfed by the hypertrophic brooding organ tissue. ^30^ The larvae are released by hatching from the egg membrane after approximately 27 days post mating (dpm; at 16°C) and remaining egg shells are discarded by the male brooding organ several hours later, when the brooding organ tissue undergoes hypotrophy.

The brooding organ of *S. typhle* evolved into a more derived, partially sealable brood pouch, which runs along the ventral side of the anterior tail Fig. 1 f). In contrast to *N. ophidion*, the brooding tissue includes two muscular skin flaps running along each side of the brooding patch parallel to the body axis, which together form a pouch when closed. Outside of the mating season, these are hypotrophic and folded flat towards each other. However, when male individuals become ready to mate, these skin flaps undergo remodeling and hypertrophy, and, together with the actual brooding patch, form a U-shaped brooding unit into which the female transfers her round eggs (∼1.5-2mm in diameter; see also ^28^). Egg numbers are smaller compared to *N. ophidion* (but still can exceed 100 in large individuals ^31^), and a filled brood pouch typically holds several dozens of eggs (depending on the male size). Occasionally, only partially filled pouches are found, possibly because mates assessed their partner’s attractiveness during mating as too low to invest the maximal possible number of eggs/pouch space into the brood.^35^ After mating, the male closes the brood pouch via the muscular skin flaps, which fold towards each other and cover the eggs completely. The inner endometrium-like skin of the brooding patch (“pseudo-placenta”) and skin flaps then engulf the eggs almost completely. The male may mate again, typically by opening the anterior part of his brooding pouch and eggs are deposited anterior to the previous clutch.^31^ *S. typhle* embryos hatch within this brooding pouch after approximately 16dpm (at 16°C) and are released after approximately 31dpm. Therefore, the primary difference between the brood structures of the two pipefishes investigated is that *N. ophidion* possesses an exposed brood patch whereas *S. typhle* provides a more shielded incubation environment offering increased opportunities for parental provisioning through interaction with an endometrium-like tissue.

The brooding organ of *H. erectus* is the most derived type found in syngnathids (Fig. 1 g). In contrast to *S. typhle*, the two skin flaps become fused at the midline during brooding organ development,^36^ forming a sack-like pouch with only a small, muscular opening at the cranial end of the tail. Pear-shaped eggs (approx. 1.5mm wide and 2.5mm long) are deposited during mating into this pouch, where they are fertilized and engulfed by the hypertrophic endometrium-like tissue. Egg numbers again depend on male and female body sizes and range from <100 to >1500 in the wild.^34^ Embryonic development commences inside the egg but embryos often hatch well before being released: we observed that in clutches of similarly developed embryos some hatched as early as 6dpm in the pouch while others had not yet hatched at 12dpm, suggesting that hatching may be determined by the microenvironment in the pouch, rather than being ontogenetically synchronous. After 16dpm juveniles were released from the paternal pouch (at 23°C), whereby some heterogeneity in juvenile developmental stages can be observed between batches.

### Pre-release development as revealed by microscopy

Pre-release development was documented using microscopy after extraction of embryos from egg-bearing males of *N. ophidion*, *S. typhle* and *H. erectus*. Table 2 gives an overview of the timed appearance of benchmark morphological features. Our description of pre-release development is intended as a comparative, yet global, overview of syngnathid ontogeny and is arguably more limited in detail concerning morphology and developmental timing than staging series published for model species (e.g. zebrafish, medaka ^37–39^). As in previous studies, the difficulties in obtaining syngnathid embryos combined with the fact that developmental speed among and within batches can vary somewhat (as reported before) made obtaining targeted stages challenging. Thus, references to developmental time should be treated as approximations (e.g., 10dpm may mean anything from 9dpm+12h to 10dpm+12h; see Experimental Procedures section for more details). Previous literature divided syngnathid pre-release (or “pre-natal”) development into varying stages,^20^ but the description provided here will follow these only loosely, and rather refer to developmental events and when these occur in the three considered species (in days post mating, or relative developmental time in percentage) as recognizing comparable developmental stages is quite subjective (as discussed in ^24^). We adhere to terminology for “direct” and “indirect” developing species as used in ^13–15^. Herein the term “larva” is reserved for the first free feeding (i.e., post-embryonic) stage, but only in cases where important differences with the adult body plan exist that only disappear upon metamorphosis, as is the case for the indirect developing *N. ophidion*. We consider *H. erectus* and *S. typhle* direct developing species in which the embryo directly transforms into a feeding juvenile, as defined by the presence of an adult-like body plan. In our description of syngnathid ontogeny we first present the overall pre-hatch/pre-release development (early, middle, late) while focusing on characteristics recognizable without further staining. Subsequently, a more detailed description of osteology is provided based on stained samples. Furthermore, *in situ* hybridization was used to explore early skeletogenesis and myogenesis.

**Table 2:**
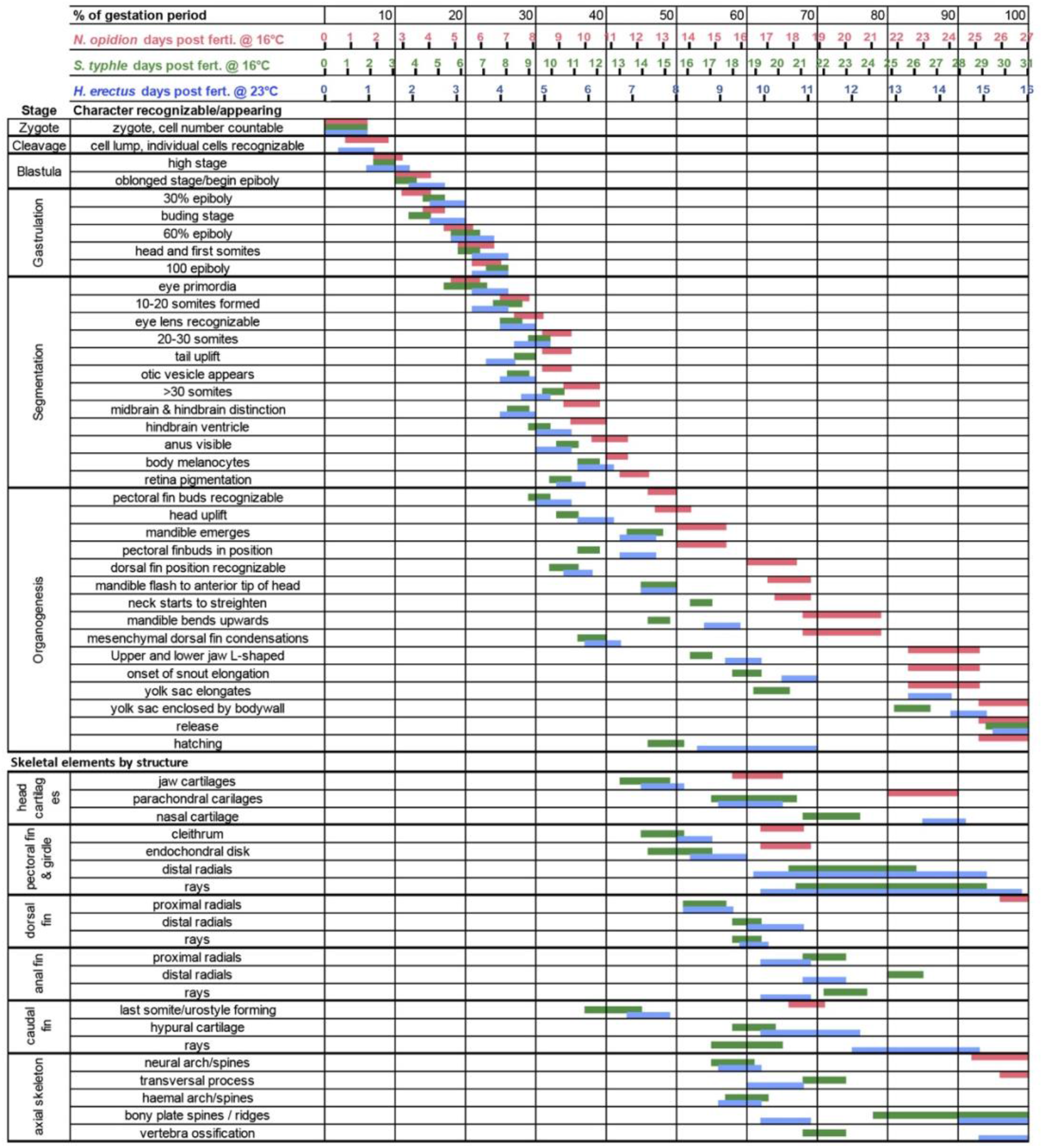
Emergence sequence of syngnathid morphological characteristic during their pre-natal development.

#### Early pre-release development: from mating to mid-hindbrain barrier

The first third of development in all three studied syngnathid species is similar and synchronized with respect to relative developmental time (Fig. 2; Table 2): in the first days after mating (zygote, blastula & segmentation stage), all embryos pass through the typical stages of early teleost development, including cleavage (with high, oblong, dome stage and onset of epiboly) and gastrulation (epiboly commences, the embryonic shield forms and the tail bud becomes visible; for reference see, for instance, zebrafish staging ^37^). Eye cups and otic vesicles emerge during the segmentation stage, which is when somites become visible. When the mid-hindbrain barrier becomes visible, *N. ophidion* embryos have completed approximately 1/3 of their pre-hatching development (9dpm), encircle the relatively small yolk sac almost entirely, and have developed over 30 somites (Figure 2 upper panel). In contrast, *S. typhle* reaches this stage somewhat earlier (8.5dpm) but with fewer than 20 somites formed (Figure 2 mid panel), similar to *H. erectus*, which reaches this stage approximately after 4dpm, i.e., 25% of its pre-release developmental time (Figure 2 lower panel). The latter species’ embryos also appear relatively smaller compared to yolk sac size due to the larger yolk sac. At this stage a heartbeat was also recognizable in all species.

**Figure 2:**
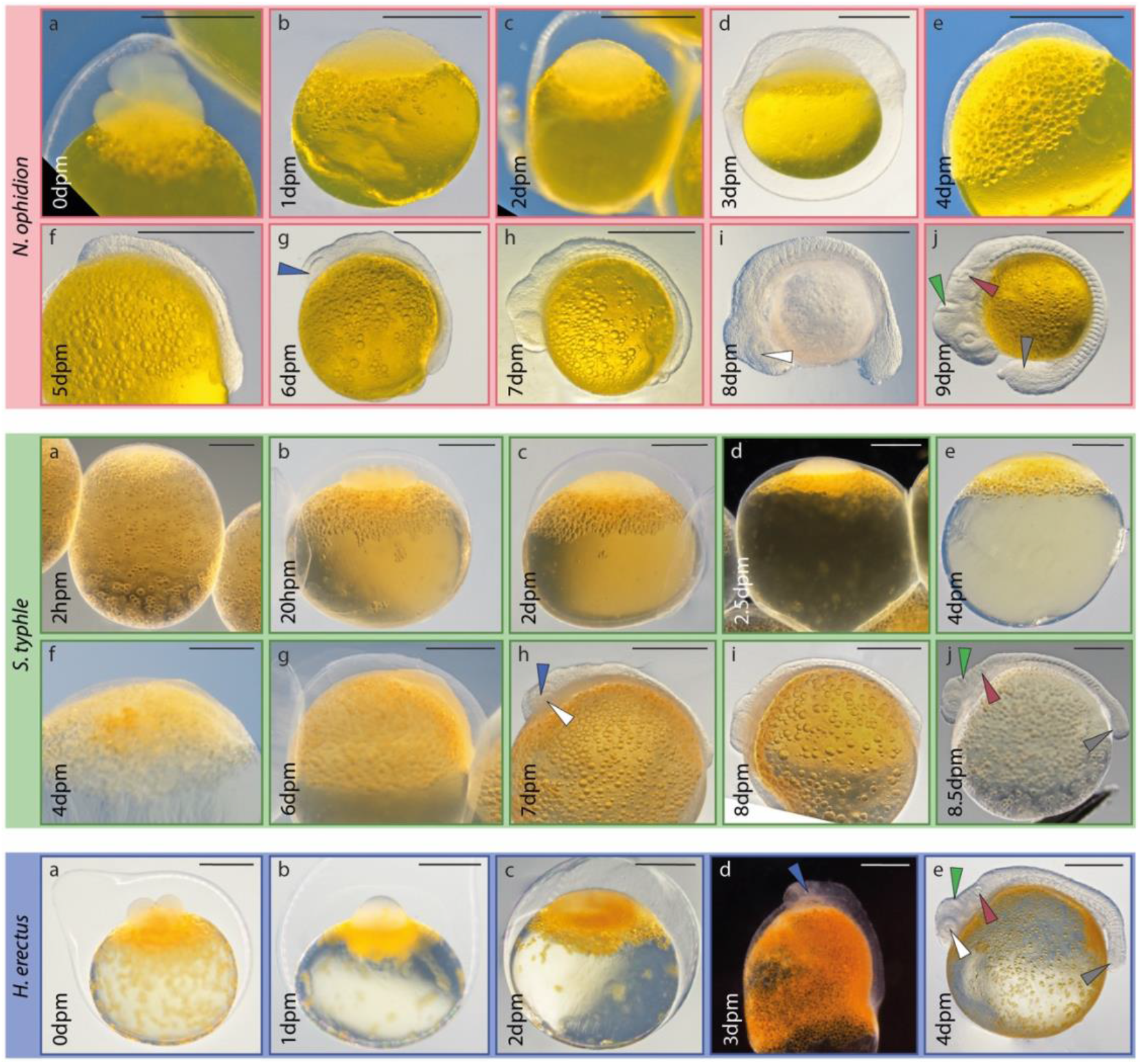
Early pre-release development of examined Syngnathids. Upper panel, mid panel and lower panel are *N. ophidion*, *S. typhle* and *H. erectus*, respectively. Overall early development is similar among studied species. Arrowheads: blue is eye cups; white is lens; red is otic vesicle; green is mid-hind-brain barrier; grey is tail bud. Scale is 500µm; dpm = days post mating.

#### Mid pre-release development: appendages and jaw emerge

After the mid-hindbrain barrier is established, the hindbrain ventricle forms visibly and soon after the first pigment cells can be detected in the dorsal part of the eye’s pigment layer and along the trunk (Fig. 3). Pigmentation continues to spread during the following days, especially along the trunk and head, and the eyes become dark and fully pigmented. Somitogenesis continues until day 18 in *N. ophidion*, which never develops a caudal fin bud, and until 12 and 7, in *S. typhle* and *H. erectus*, respectively, after which caudal fin buds form. Mandibular condensations are visible at approximately 15dpm, 10dpm and 8dpm in *N. ophidion*, *S. typhle* and *H. erectus*, respectively, which continuously grow and extend anteriorly. Dorsal fin condensations can be identified after 10dpm and 6dpm in *S. typhle* and *H. erectus*, respectively, while this takes until 17dpm in *N. ophidion*. The latter species is characterized by a prominent fin fold along the caudal region, which starts to form at 15-17dpm and is resorbed in the second and third week after hatching. In *H. erectus*, during this period (starting at ∼6dpm) the hindgut is noticeably increasing in volume, a condition maintained after release.

**Figure 3:**
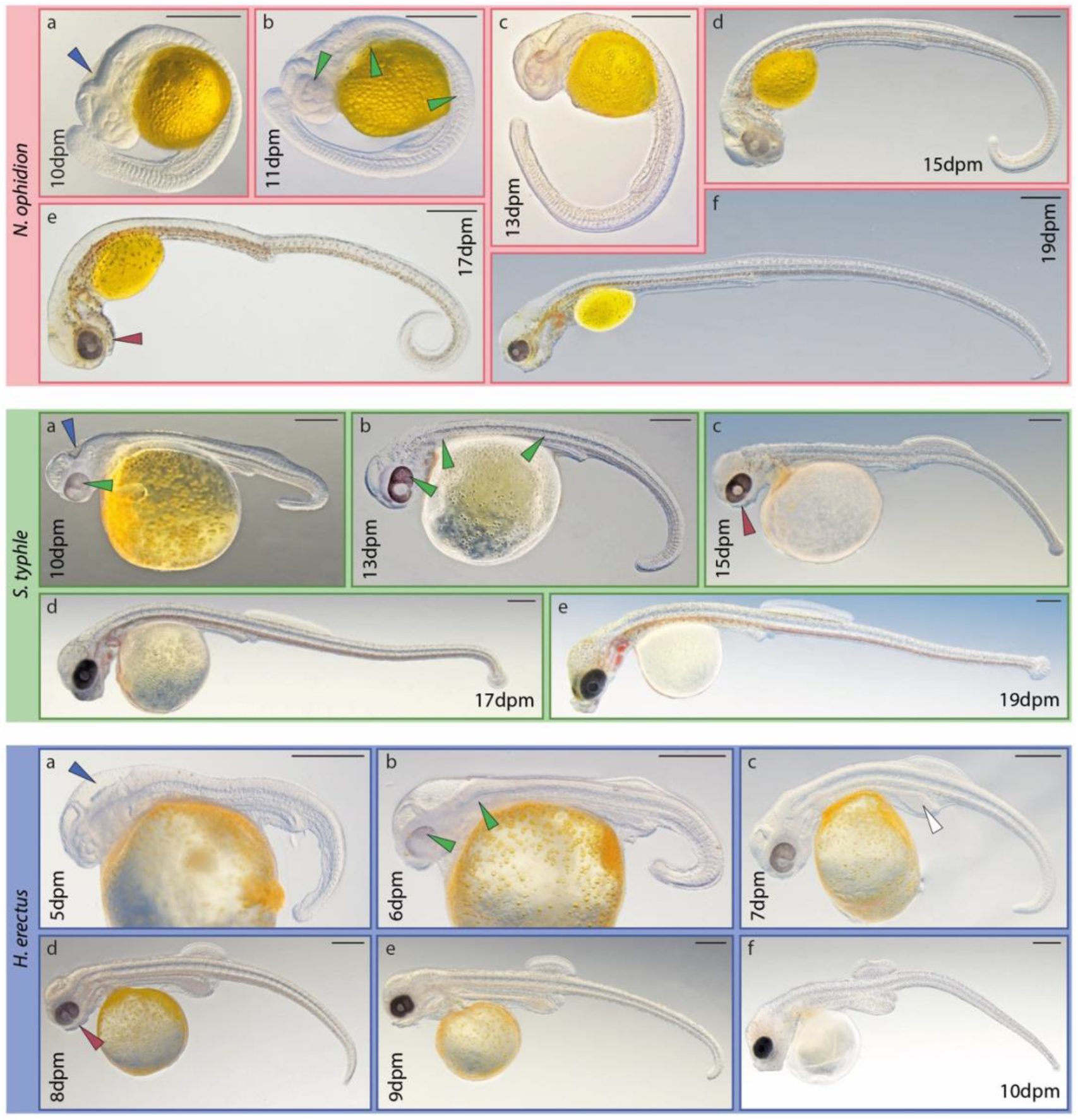
Mid pre-release development of examined syngnathids. Upper panel, mid panel and lower panel are *N. ophidion*, *S. typhle* and *H. erectus*, respectively. In this period species-specific characteristics develop more clearly. Arrowheads: blue is hind brain vesicle; green is first pigmentation; red is the lower jaw; white is the hind-gut hypertrophy. Scale is 500µm; dpm = days post mating.

#### Late pre-release development: jaw outgrowth, yolk resorption, bony plates emergence

The last third of pre-release development is characterized by increased pigmentation, yolk sack resorption, the elongation of the tube-shaped mouth, and the appearance of the characteristic bony plates in *S. typhle* and *H. erectus* (Fig. 4). While *S. typhle* and *H. erectus* become fully pigmented (except for some fins) during this period, *N. ophidion*’s pigmentation is still patchy at the day of hatching and full pigmentation cover of the body is only achieved within the first week post hatching. In all three species the yolk sack is typically fully absorbed when juveniles are released by their fathers. Interestingly, both pipefishes pass through a stage during which their head trunk connection resembles a seahorse. This is most striking in *S. typhle* (which is more closely related to seahorses than it is to *N. ophidion*). From 21 dpm to 28 dpm the head and trunk are positioned at a right angle from each other, which becomes only parallel towards release (31dpm). In contrast in *H. erectus* the initial right angle between head and trunk never straightens but is maintained in adults, possibly indicative of developmental truncation (see discussion).

**Figure 4:**
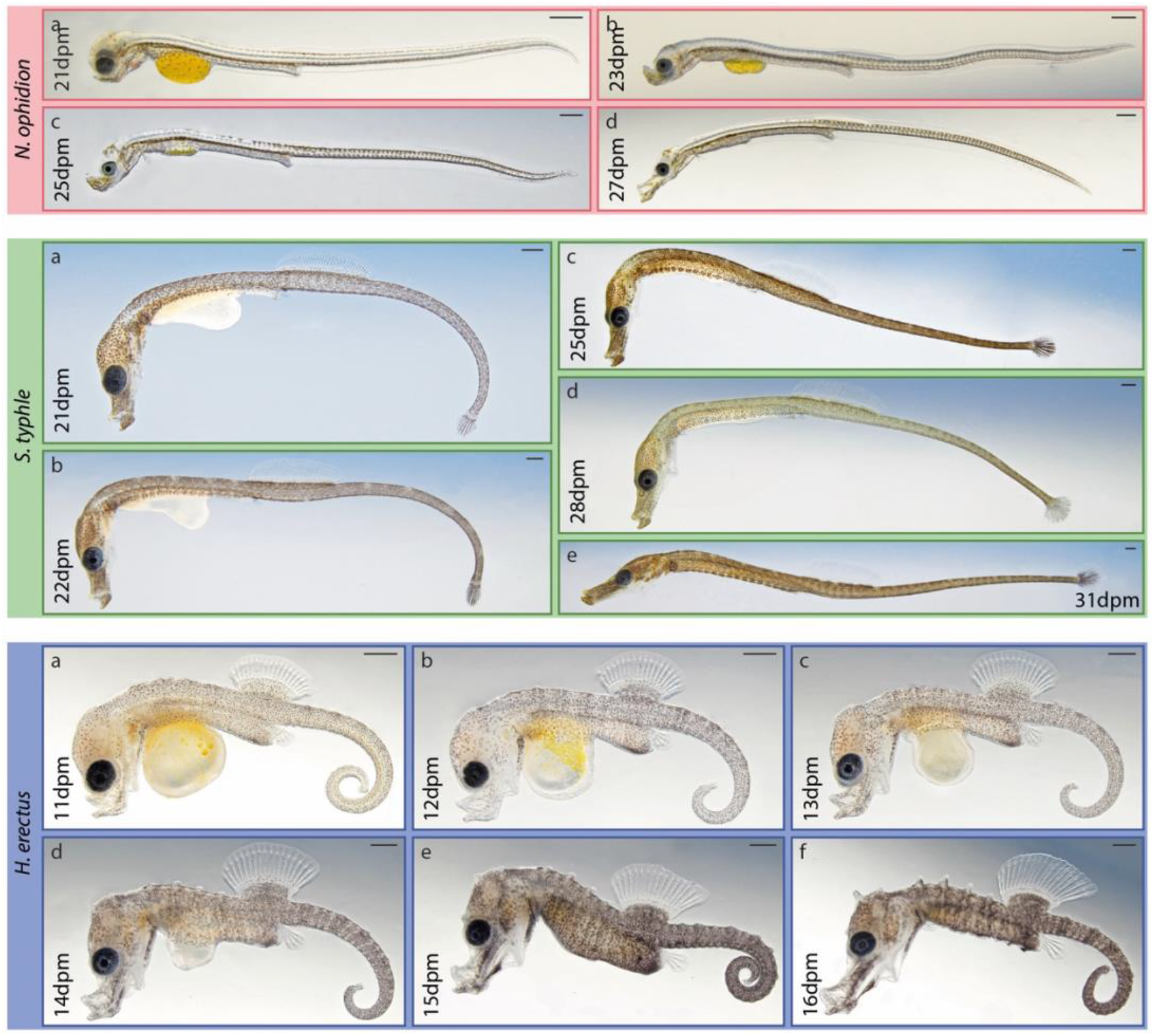
Late pre-release development of examined syngnathids. Upper panel, mid panel and lower panel are *N. ophidion*, *S. typhle* and *H. erectus*, respectively. The last pre-release period is characterized by snout elongation, continued pigmentation and the conclusion of allometric fin outgrowth. Scale is 500µm; dpm = days post mating.

### Skeletal development as revealed by Alcian blue and Alizarin red staining

The skeletal development of studied species is presented as revealed by Alcian Blue (stains cartilaginous tissues blue) and Alizarin Red staining (stains ossified tissues red). As delicate ossification appeared undetectable via Alizarin Red staining when used as part of the low-acid (and even non-acid) double staining technique in conjunction with Alcian Blue in this study ^40^, additional Alizarin Red staining on selected stages were used to describe ossification patterns separately (see also Experimental Procedures section). This has also allowed us to detect ossifying tissues earlier than previous studies.^18^ A schematic of a *S. typhle* juvenile (chondro)skeleton is used to visualize the position of presented body features in the figures. The branchial basket is not included in the description as the used staining techniques did not reveal its structures sufficiently. Furthermore, individual structures (and substructures) may not be described if they could not be identified with the applied methods. For complementary descriptions of the (cranial) cartilages and bones in syngnathids, see ^9,10,19^.

#### Chondrocranium: the formation of a tube-shaped snout

One of the most iconic shared features of all syngnathids is their snout-like jaw apparatus, which forms the anterior part of a distinct head (Fig. 6-10). The posterior part of the head is connected to the trunk by a neck region that allows for some individualized movement (particularly in the dorsoventral dimension) of the head (depending on the species). The chondrocranium develops throughout the second half and the latter two thirds of pre-release development in the *N. ophidion* and *S. typhle* & *H. erectus*, respectively, and the only structures distinctly stained before were the otoliths/otic cartilage. Using our staining techniques, the earliest non-otic chondric condensations could be identified as the anterior tip of Meckel’s cartilage, which scaffolds the dentary (identifiable at approx. 16, 13 and 7dpm, in *N. ophidion*, *S. typhle* and *H. erectus*, respectively). Most major cranial cartilage condensations could be identified only one to two days later in all examined species, including the ethmoid plate cartilage, the palatoquadrate, the hyosymplectic cartilage, the ceratohyal and interhyal, and the parachordal cartilage. At approximately 20, 17 and 10dpm the basihyal, hypohyal and pterygoid process of the quadrate are recognizable in *N. ophidion*, *S. typhle* and *H. erectus*, respectively. The mandible (lower jaw) becomes more distinct and some days later starts to bend upwards (dorsally), forming a near vertical orientation of the mouth opening. At the same time the upper jaw emerges. While its dorso-ventrally aligned jaw position develops after 17dpm and 10dpm (corresponding to ∼55% and 63% of pre-natal developmental time) in *S. typhle* and *H. erectus*, this stage of jaw development is reached only after 20dpm (74% of pre-natal developmental time) in *N. ophidion* (Fig. 10). Interestingly, at this stage the developing ethmoid cartilage is not straight, but rather has a curved shape, which straightens during the last stages of pre-release development. This is the stage when the iconic snouts forms, starting at about 25dpm in *N. ophidion*, 19dpm in *S. typhle* and at about 12dpm in *H. erectus*. Also, during this time the vomer cartilage/rostral cartilage emerges, which is accompanied by an overall elongation of the ethmoid and hyosymplectic cartilage in combination with a re-orientation of the palatoquatrate, which rotates from its orientation rather parallel to the hyosymplectic cartilage to an orientation almost perpendicular to it. Allometric snout elongation continues after release, however, the composition and relative orientation of morphological structures to each other does not change anymore considerably.

**Figure 5:**
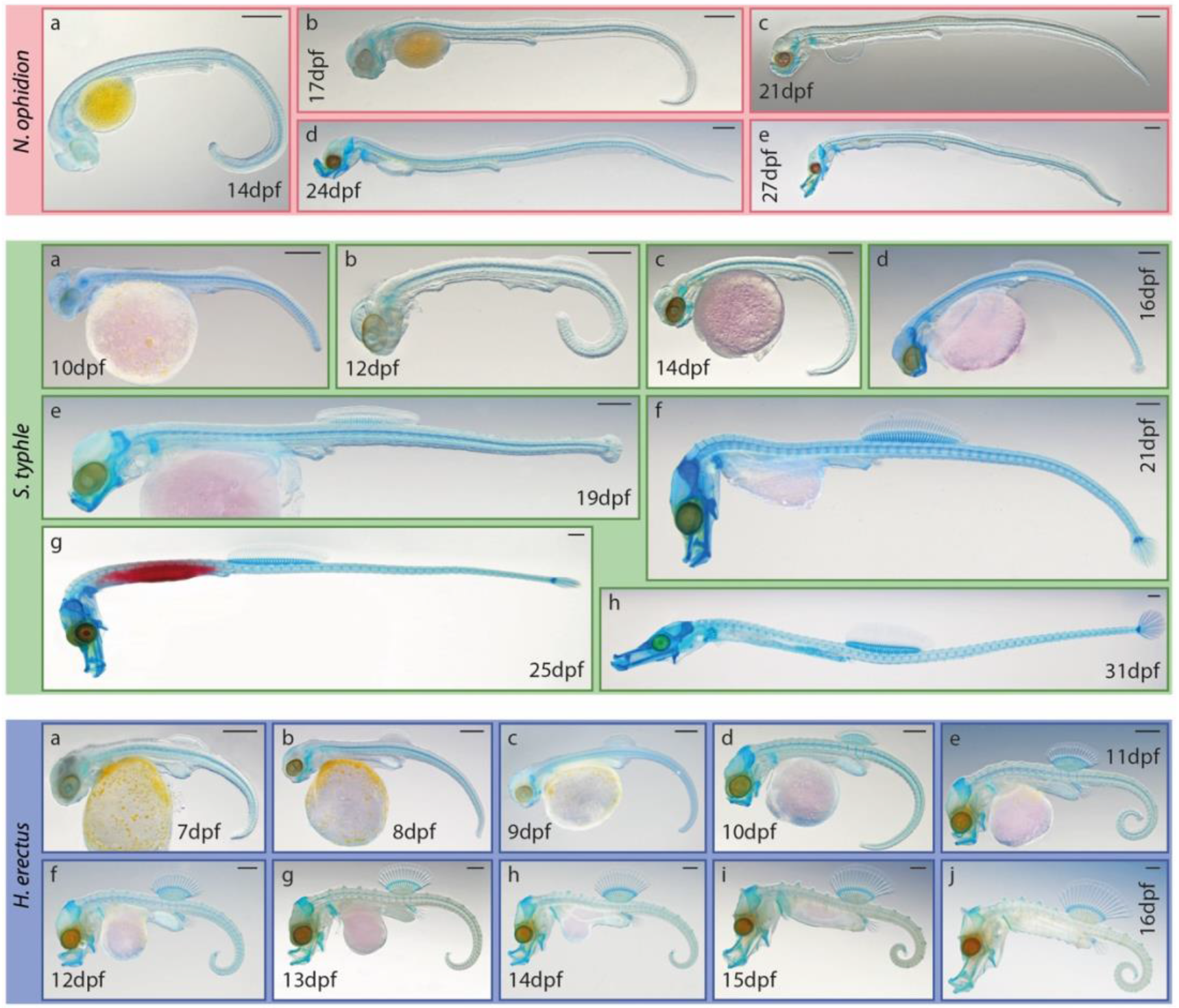
Overview of chondrogenesis across investigated syngnathids pre-natal development (Alcian Blue stained specimens). Scale is 500µm; dpm = days post mating. For red yolk coloration in mid-panel (g) see Experimental Procedures.

**Figure 6:**
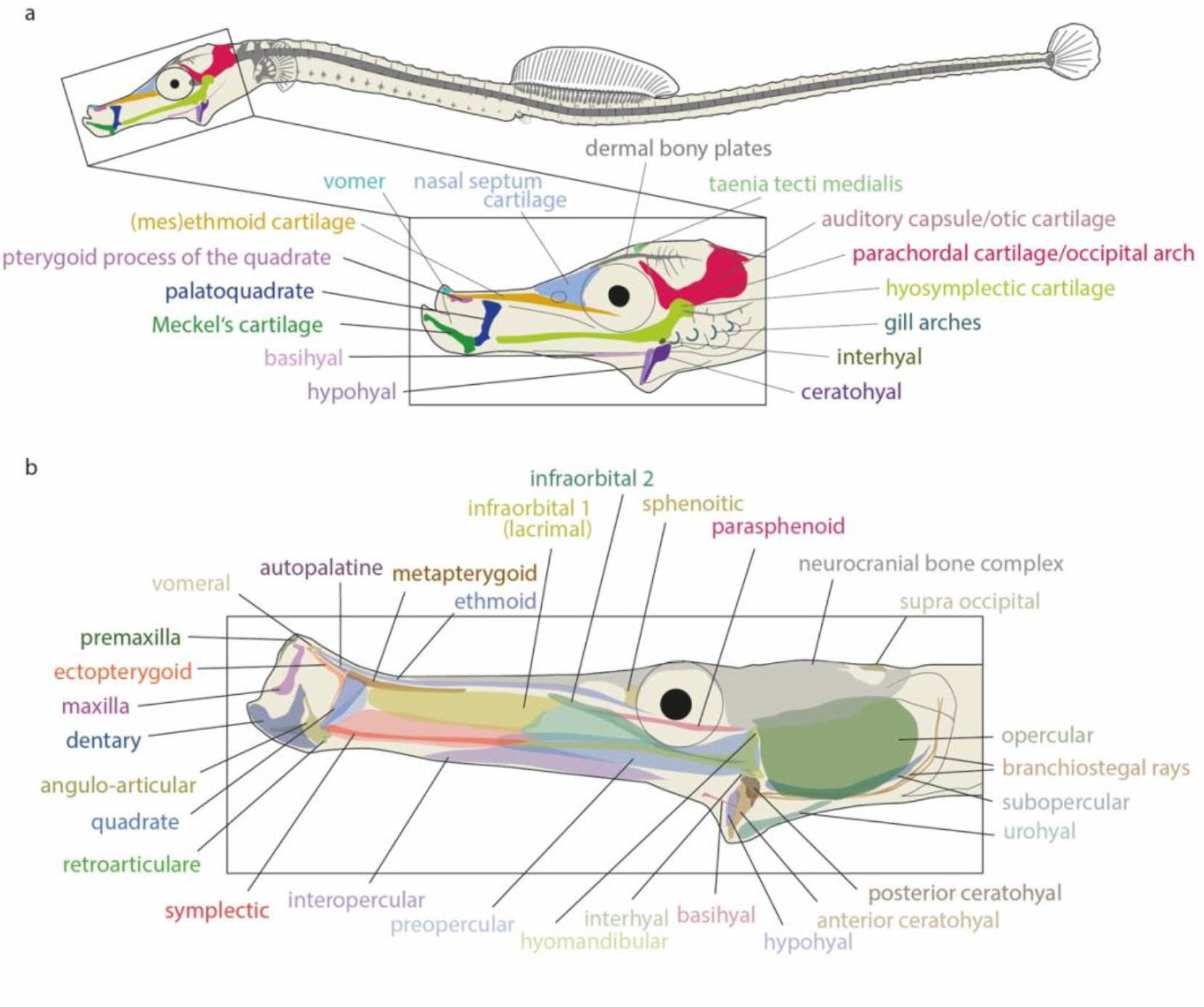
Generalized overview of the head skeletons of syngnathids. (a) Head cartilages in syngnathid embryos identifiable after Alcian Blue staining in this study. The branchial basket is not detailed. (b) Bones identifiable in adult syngnathids after Alizarin Red staining in this study.

**Figure 7:**
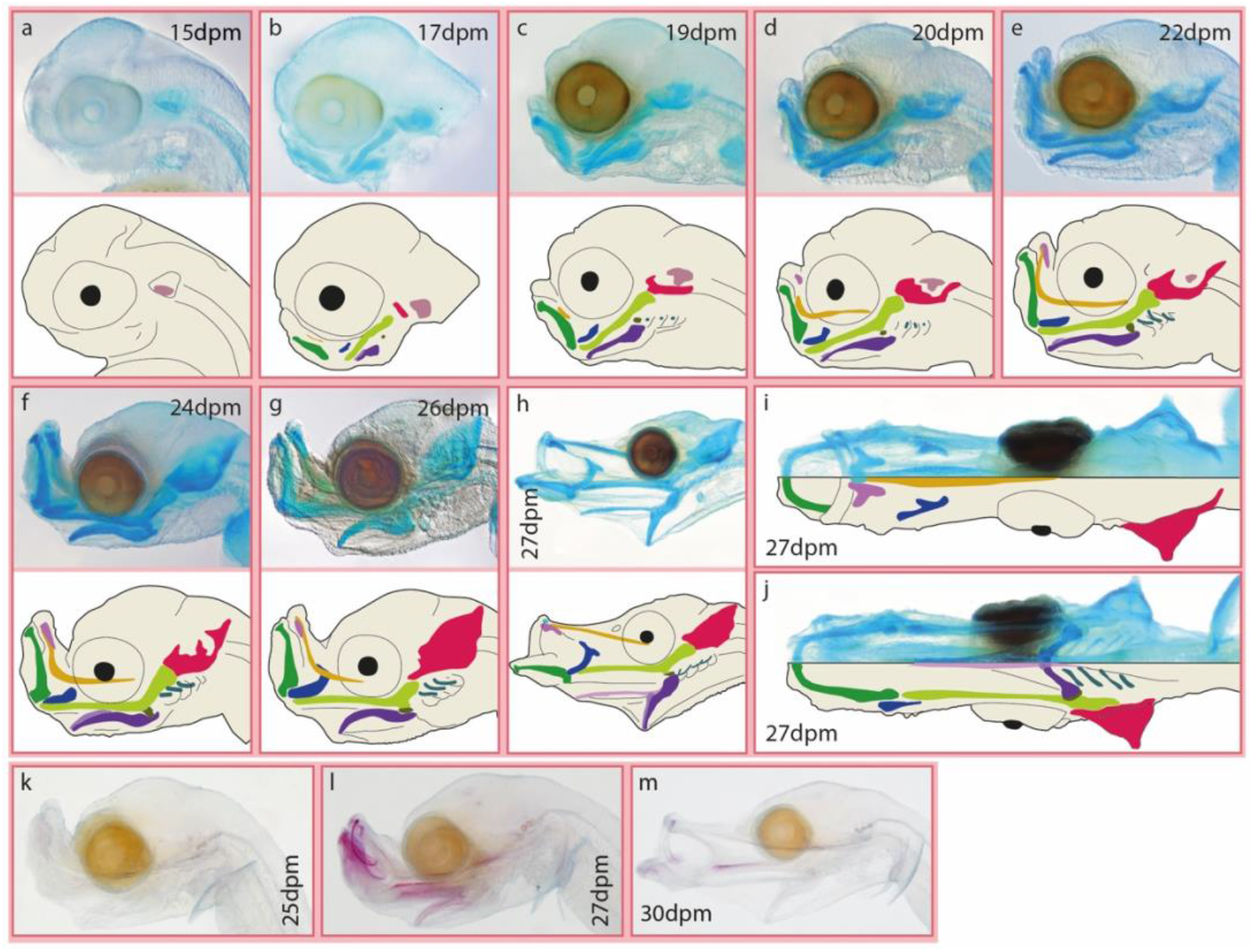
Head skeleton of pre-release *N. ophidion*. (a-j) Head cartilages and (k-m) ossifying tissues across the pre-natal development; (i) dorsal view, (j) ventral view, all others lateral view; dpm = days post mating.

**Figure 8:**
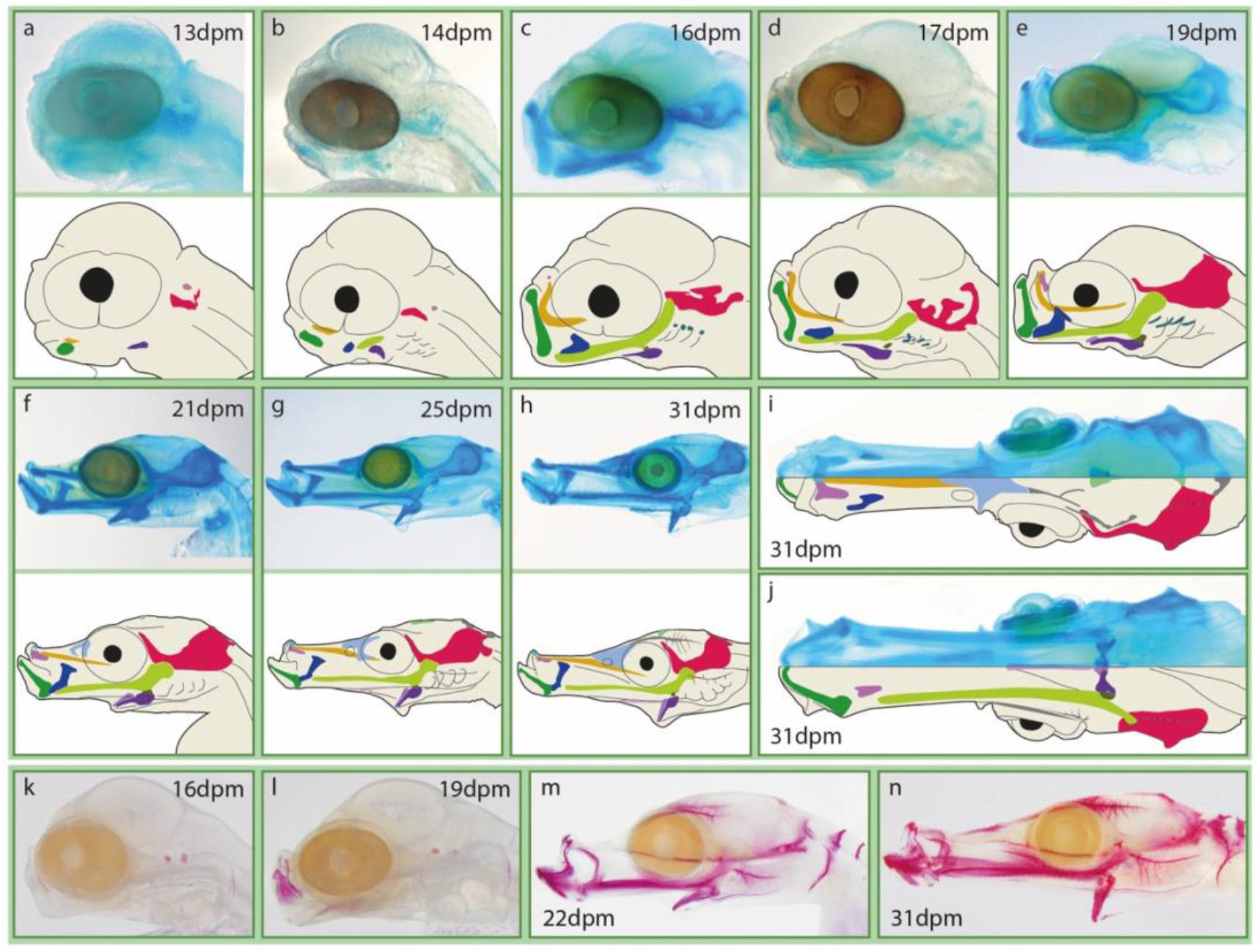
Head skeleton of pre-release *S. typhle*. (a-j) Head cartilages and (k-n) ossifying tissues across the pre-natal development; (i) dorsal view, (j) ventral view, all others lateral view; dpm = days post mating.

**Figure 9:**
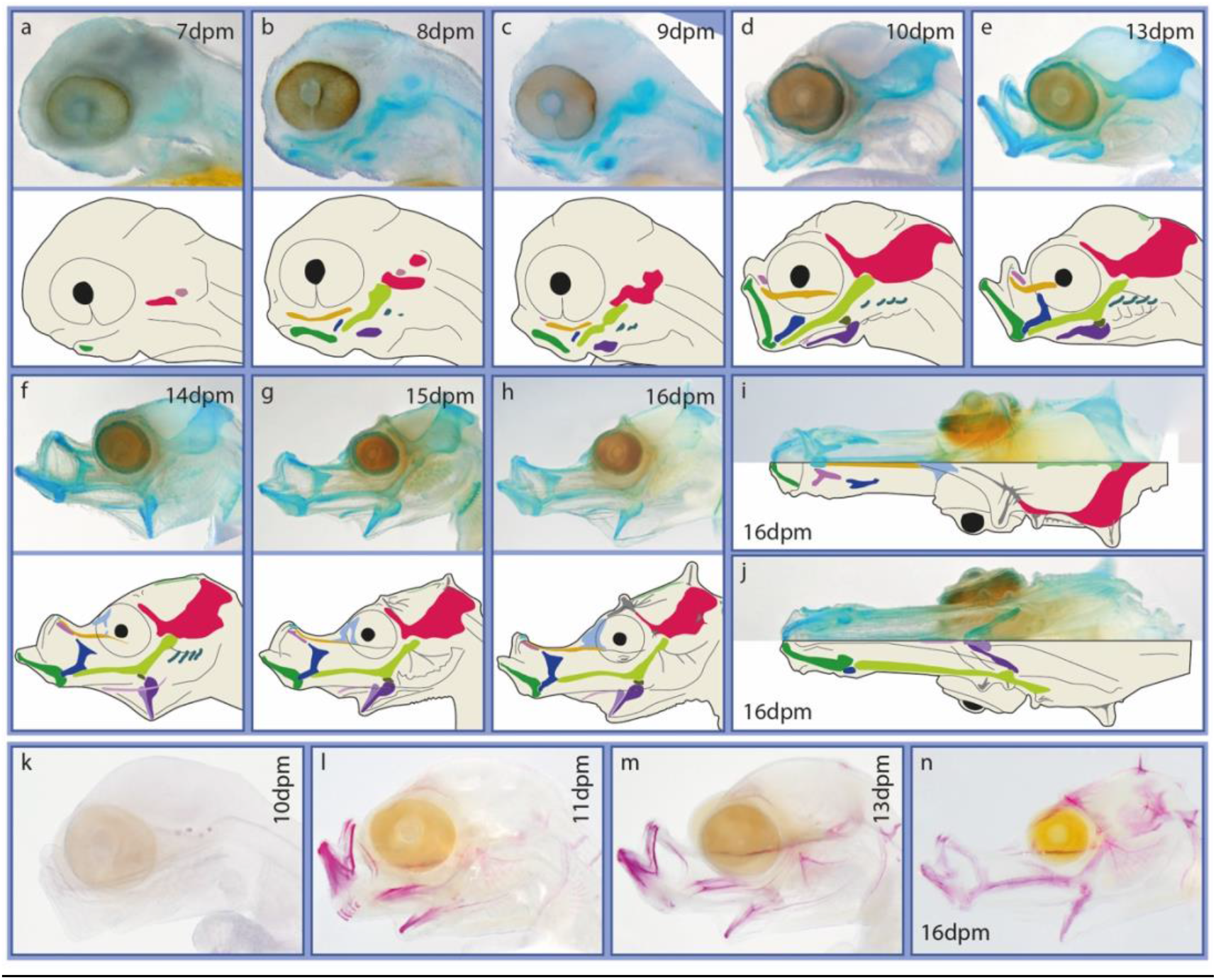
Head skeleton of pre-release *H. erectus*. (a-j) Head cartilages and (k-m) ossifying tissues across the pre-natal development; (i) dorsal view, (j) ventral view, all others lateral view; dpm = days post mating.

**Figure 10:**
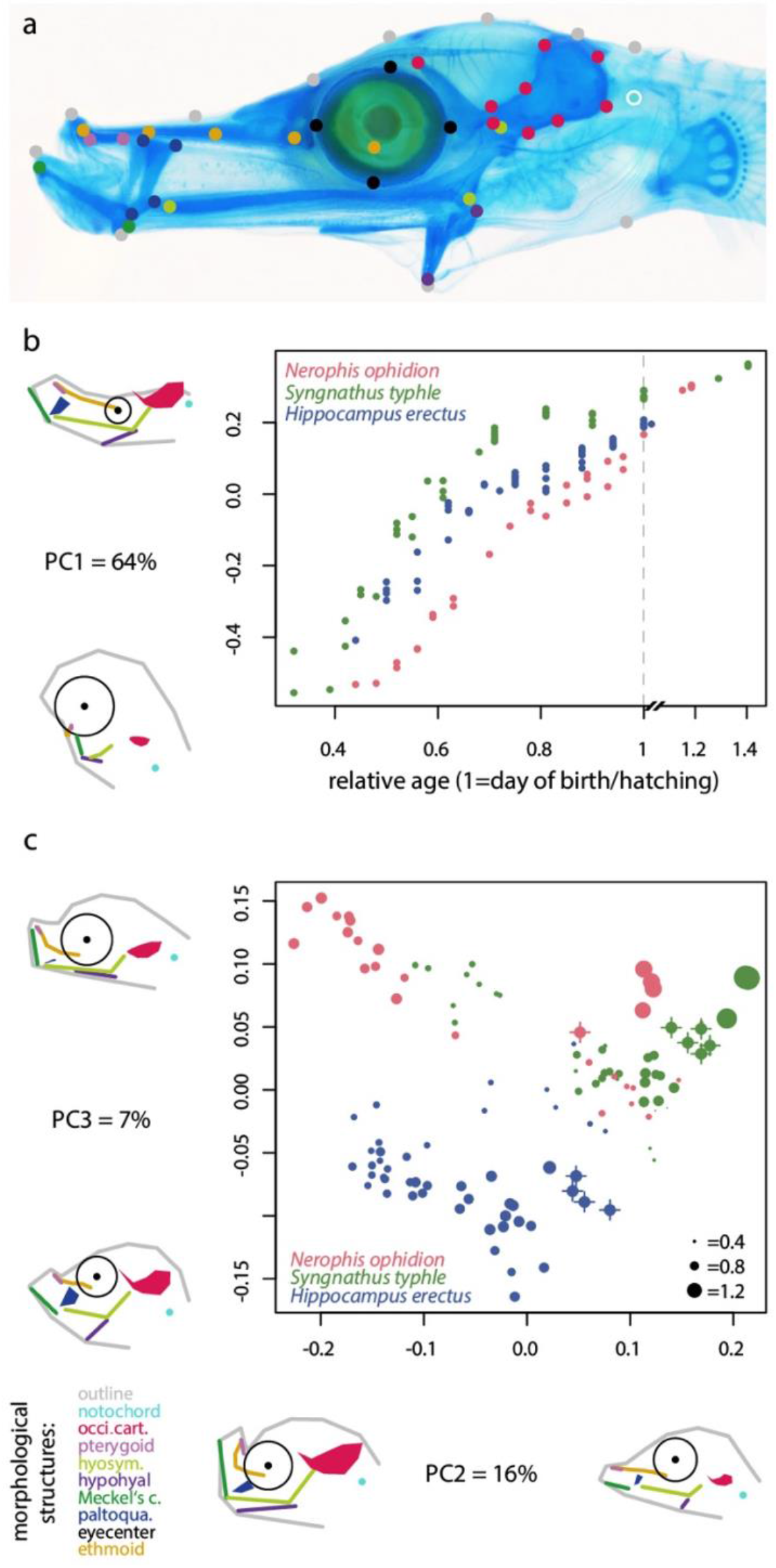
Comparative geometric morphometric PCA analysis of overall head shape and key cartilages across early development. (a) 44 landmarks were used to describe the head shape and key cartilages. In younger specimens the location of landmarks was estimated as the PCA analysis required a full dataset across all investigated stages. (b) Principal component 1 scores plotted against relative developmental age (i.e., 0=day of mating, 1=day of birth/hatching) illustrate dominant morphological variance associated with relative age/size. (c) PC2 and PC3 scores illustrate divergent developmental trajectories for the three species. Head shapes illustrate the landmark configurations at the extremes of the respective axis. Dots with crossing lines in (c) indicate release/hatching stage and dot size reflects relative age.

Geometric morphometric data on the head shape and key cartilages was collected and analyzed using PCA (see Experimental Procedure section for details and limitations of this approach; Fig. 10). Principal component one (PC1) positively correlates with relative age of the samples, suggesting it reflects the main morphological transitions investigated syngnathids’ heads underwent (Fig. 10 b). At release (rel. age =1), *S. typhle* individuals show the highest scores (i.e., elongated snout, shallow head) when compared to the other two species, while *N. ophidion* only catches up to *S. typhle* after hatching, and *H. erectus*’ scores remain intermediate between the other two species until release. *H. erectus* develops its characteristic high head cranium, which differentiates this species from the pipefish species as reflected on PC3 (Fig. 10 c). PC2 illustrates how *N. ophidion* undergoes a more pronounced L-shaped jaw stage compared to *S. typhle* (with more negative values on PC2), and only around the day of hatching the snout straightens and a morphology more similar to that of *S. typhle* is developed (i.e. more positive scores on PC2; see also Fig. 11 a,b).

**Figure 11:**
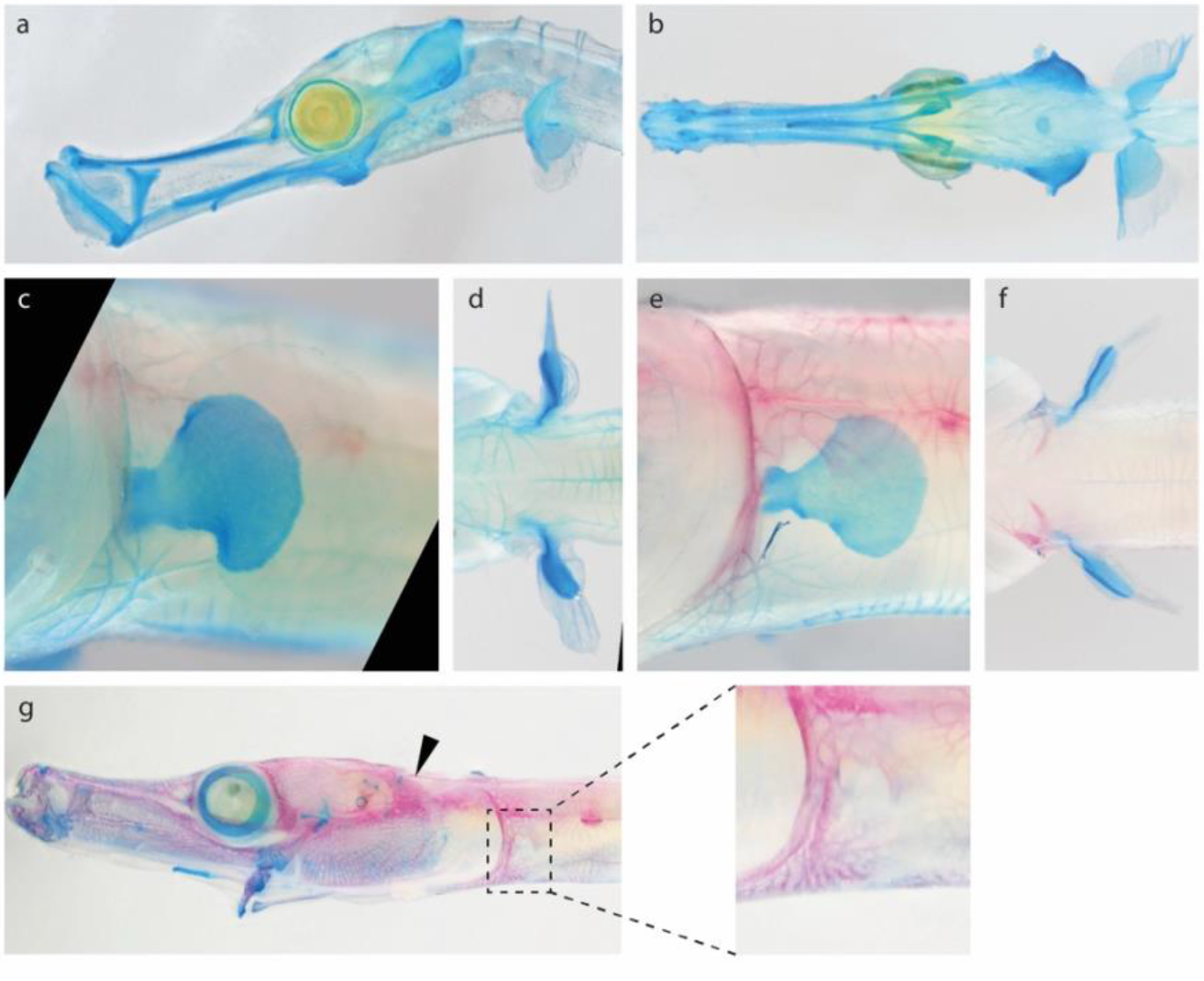
*N. ophidion* cranial and pectoral development post hatch. a,c,e,g = lateral views; b,d,f = ventral view; a,b = 10 days post hatch, c-g = later subadult stages, wild-caught individuals. Black arrowhead indicates the epaxial sesamoid bone present in *N. ophidion*.

Using separate Alizarin Red stainings, ossification was detected at earlier stages compared to previous studies,^18^ and individual bones – especially in the complex head skeleton – could be identified, although individual bones contributing to the neurocranium could not be reliably delineated (Fig. 6-10). In all studied species, otoliths were the first structures showing the onset of ossification with this technique, followed by the maxillary and cleithrum (25, 16, 10dpm, for *N. ophidion*, *S. typhle* and *H. erectus*, respectively) and shortly after the premaxillary, symplectic and dentary (Fig. 7-9). In *N. ophidion*, Alizarin Red staining suggests that ossification indeed does not commence much further until the larvae hatch and only few more structures show staining, such as the parasphenoid, ceratohyoid, and basioccipital (Fig. 7 lower panel). In *S. typhle* and *H. erectus*, ossification commences much further until the day of release and then can be detected in most facial/cranial bones (Fig. 8,9 lower panels).

#### Formation of the occipital and pre-caudal axial skeleton

In vertebrates the somites contribute to all the body regions except for the anterior head. Four main somite derived skeletal regions are distinguished in fishes, namely the basioccipital (posterior part of head region), precaudal (trunk), caudal (tail) and ural (caudal fin) region.^13^ In teleosts, somites 1 and 2 contribute to the basioccipital, which forms the posterior part of the cranium, and somite 3 connects the cranium to somite 4, which represents the first somite developing into a vertebra.^13^ We examined the formation of the axial skeleton using Alcian blue and Alizarin red staining. In all three examined syngnathids, the notochord is visible as a subtle blue rod already at very early stages (e.g. in *S. typhle* at 8dpm; Fig. 12). The first condensations of the neural arches were observed at the most anterior precaudal somites at 24, 17 and 10 dpm for *N. ophidion*, *S. typhle* and *H. erectus*, respectively (Fig 12). It is noteworthy that, whereas in *N. ophidion* three days later neural arch formation was still restricted to the most anterior somites, neural arch protrusions were observed in all precaudal and most caudal somites in *S. typhle* and *H. erectus* only one day after their first appearance anteriorly. In *H. erectus* and especially *S. typhle*, the neural arch elements from somites 4 to 6, which correspond to vertebrae 1 to 3, become notably hypertrophic and distinct from other neural arch elements during pre-release development. The first transversal processes were recognizable on central precaudal and anterior caudal vertebrae in *S. typhle* at around 22 dpm and in *H. erectus* at 10dpm, while in *N. ophidion* these processes could not unambiguously be identified before hatching day. The formation of the vertebral centra is not detected using Alcian blue because in teleosts these elements form through intramembranous ossification ^13^ and ossifying vertebrae centra are observed via Alizarin Red staining at 22dpm and 16dpm in *S. typhle* and *H. erectus* (Fig. 12 lower panel), while no ossification of vertebrae centra could be detected in *N. ophidion* before hatching. In contrast, in *S. typhle* and *H. erectus* the neural & haemal arches/spines, transversal processes, dorsal and anal fin radials and rays, caudal fin rays, and pectoral proximal fin radials already undergo ossification at the day of release according to Alizarin Red staining. Interestingly, Alcian blue stains the non-ossifying part of the vertebral column (Fig. 12), which probably corresponds to the intervertebral disks.

**Figure 12:**
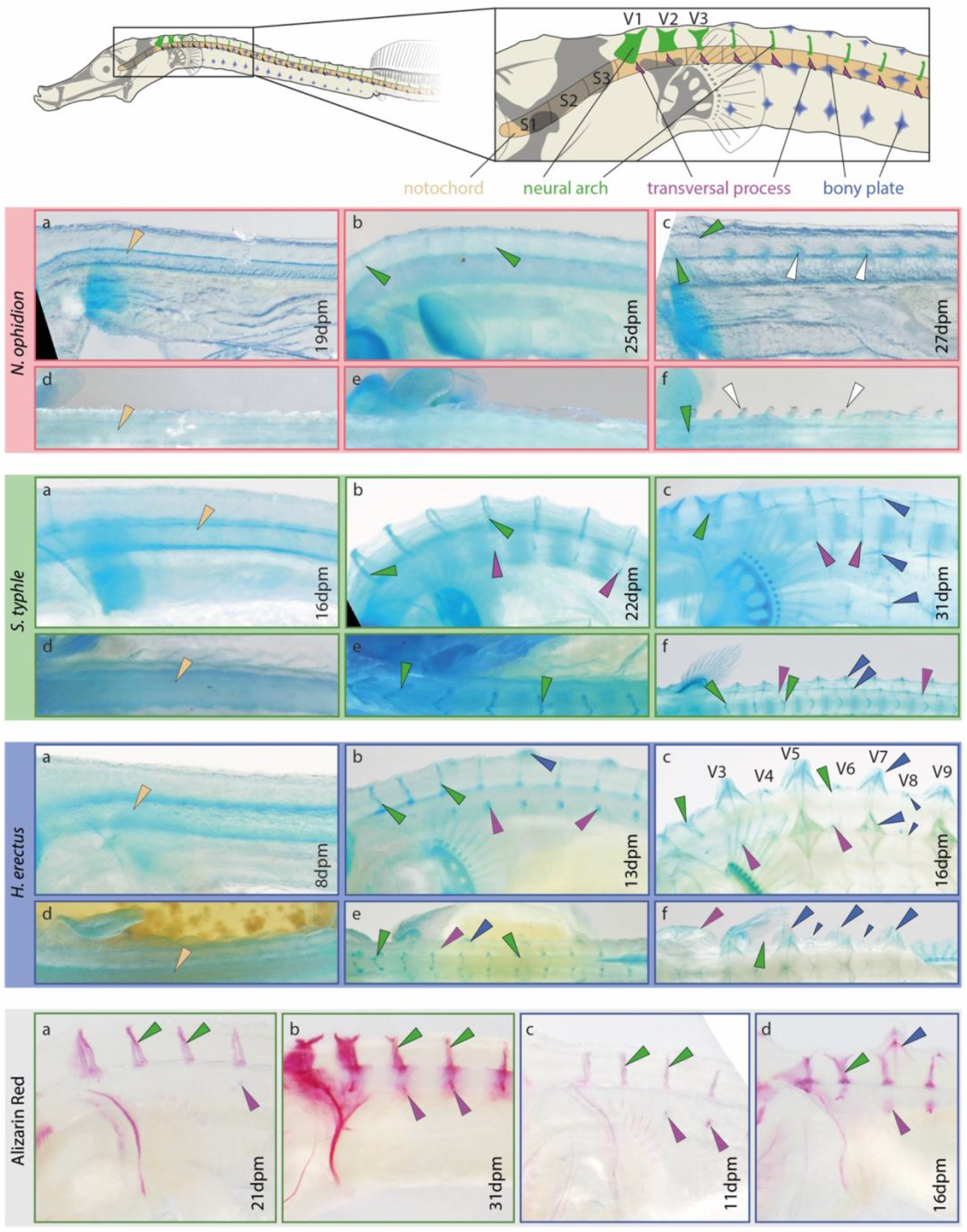
Development of the syngnathids’ axial skeleton. Upper panel, second and fourth panel: main skeletal structures contributing to the anterior axial skeleton in *S. typhle*, *N. ophidion* and *H. erectus*, respectively. White arrow heads indictae spine-like structures (not prominently present in *S. typhle*). (a-c) Lateral view, (d-f) dorsal view. Lower panel: ossifying tissues of the anterior axial skeleton across the pre-natal development; all lateral view; (a,b) *S. typhle*, (c,d) *H. erectus*; dpm = days post mating.

Ribs and epicentrals could not be identified in any of the studied species in line with the notion that syngnathids have secondarily lost ribs,^1^ and sesamoid bones in epaxial tendons were only identified in subadult or older stages of *N. ophidion*, but not in *S. typhle* nor *H. erectus*, consistent with previous studies (Fig. 11).^10,41^ The emergence of the haemal arches, which define the caudal region is discussed below together with the developmental of the caudal region.

#### Development of the syngnathids’ bony plate armour

Adult syngnathids possess an armour of jointed bony plates in their integument,^42,43^ which provides protection and structural support for the fins (Fig. 1 h).^12,44^ All three studied species feature a very similar pattern of seven longitudinal (i.e. anterior-posterior) oriented rows of bony plates in the neck and trunk region. Three rows cover each flank in a bilaterally symmetrical pattern whereby the dorsal most rows on each side close in on each other in the dorsal midline. The ventral side is covered in a single unpaired row of plates (Fig. 13), which is discontinued at the anus. In a transition region (approx. from anus to the posterior end of the dorsal fin) the three pairs of flank rows are reduced to the two pairs that form the majority of the square-shaped plate armour of the tail region (with one row forming each edge, Fig. 13).^42,44^ Interestingly, this transition from six to four rows is different in each species: in *N. ophidion*, the two most dorsal pairs of rows continue seamlessly into the tail region; in *S. typhle*, instead, the most dorsal pair of rows is discontinued after the posterior border of the dorsal fin and the remaining two rows form the tail armour; and in *H. erectus* only the middle trunk row is continued as the ventral tail row, while the dorsal tail row emerges in between the two most dorsal trunk rows at the eleventh plate ring (Fig. 13).

**Figure 13:**
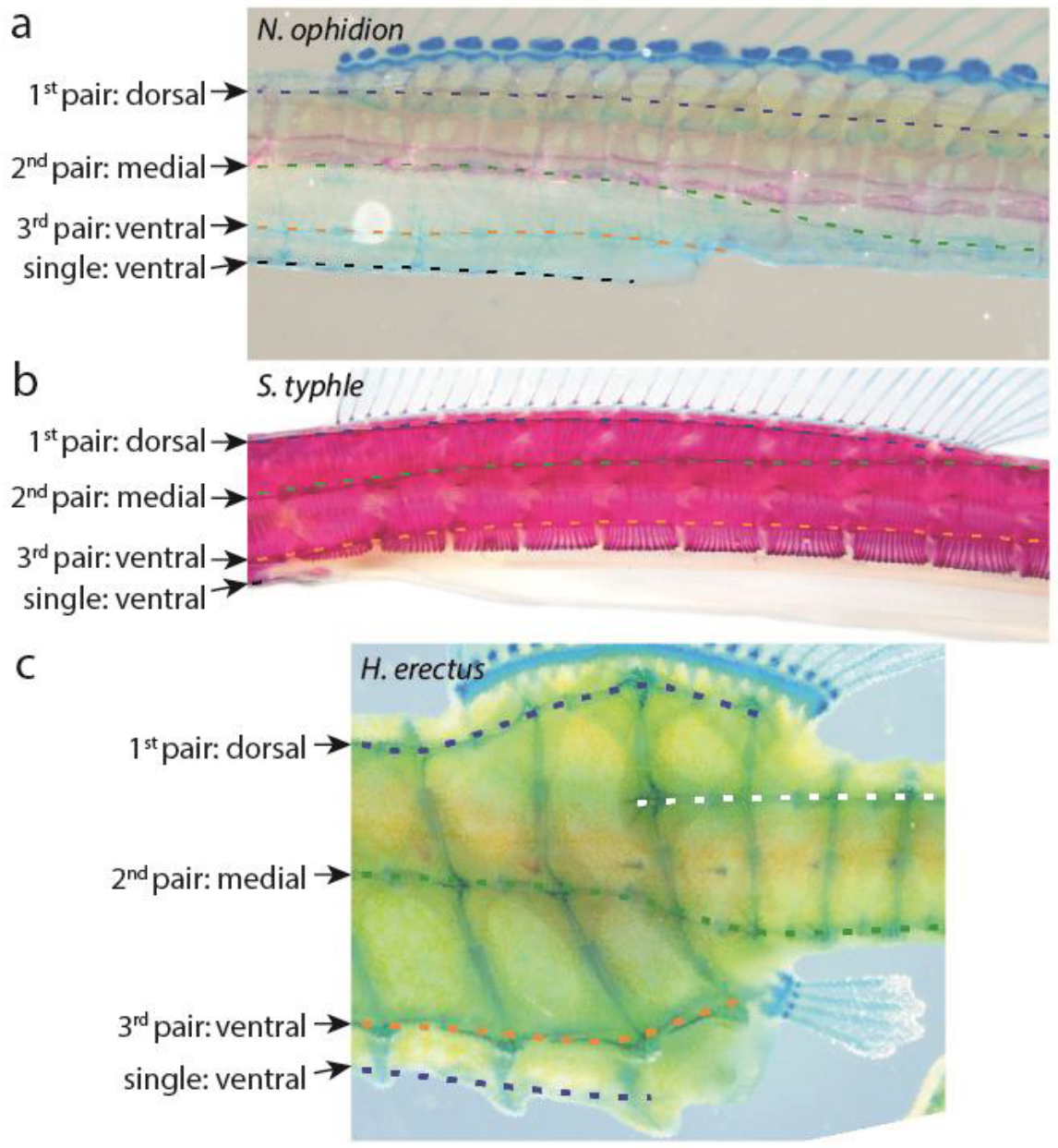
Bony plate transition from pre-caudal to caudal. The transition of the seven rows of bony plates on the pre-caudal region n into four rows on in the caudal region differs among species. *N. ophidion* (top), *S. typhle* (mid) and *H. erectus* (lower) from a lateral view (cranially left, dorsal up); individuals stained with Alcian Blue and/or Alizarin Red.

The first putative indications of integument covering in *N. ophidion* occur during the same stages the axial skeleton emerges (Fig. 12) and can be observed as one lateral row of pointy spines using Alcian blue (26dpm; Fig. 12, second panel, b,e) (which here probably is indicative of dermally ossifying condensations with high glycosaminoglycan content, rather than of cartilage). Together with spines formed by cranial cartilages (see Fig. 6 i,j) *N. ophidion* larvae exhibit thus a row of defensive spines along the body, which is only resorbed one to two weeks after hatching and likely functionally replaced by the adult bony plate armour. It remains unclear if these spines are developmental precursors of bony plates or have an independent origin. Notably, similar protrusions are also observed in *H. erectus*, starting at ∼12dpm (Fig. 12 lower panel, b). Here, protrusions remain blunt and become bony plates centers. Given the apparently divergent ontogenetic trajectories the homology between the early dermal protrusions in *N. ophidion* and *H. erectus* remains for now unclear.

Adult *N. ophidion* and *S. typhle* have similar plate types and individual plates are connected by well-developed longitudinal spine-and-groove joints and smaller (or absent) transversal peg-and-groove joints (Fig. 14a-d; for details on plate ultrastructure see ^43^). Their bony plates develop from cross-shaped tissue condensations (“ridges”) that stained blue with Alcian Blue (Fig. 10 mid and lower panel, c,f; observable from 10 days post hatch and 28dpm onwards in *N. ophidion* and *S. typhle*, respectively). These crosses emerge along the body in aforementioned rows, with one cross being formed per vertebra and row (observable from 10days post hatch and 28dpm onwards in *N. ophidion* and *S. typhle*, respectively). Crosses then continue to form the main ossification ridge of maturing bony plates throughout further development as they expand, ossify and get into contact with each other (Fig. 14). When forming bony plates start to overlap peg-and-groove sliding joints are formed with the more cranially-oriented margin of a plate always sliding over its neighboring plate and, by doing so, forming the groove. The more caudally-oriented margin slides under its neighbor and forms the peg (Fig. 14 a-d). In addition, a plate’s dorsal margin slides over its more dorsally situated neighbor, and one or multiple joints may be formed. In *H. erectus*, plates are connected by exactly one well developed peg-and-groove joint to each adjacent plate (Fig. 11 e-h). In studied pipefish, only longitudinally oriented joints (i.e., joints to cranially or caudally adjacent plates) form such well-developed joints, while those between dorsoventrally overlapping plates remain often much less well developed: here several smaller joints can be found (especially in the precaudal region) or joints can be absent (especially in the caudal region), suggesting different evolutionary demands for skeletal flexibility among species and body regions within species.^42,44^ Furthermore, the shape of these plates in studied pipefishes could be visualized using Alzarin Red staining only relatively late in development (for both pipefish species, weeks after hatching/birth), suggesting that rigidity of the bony plate armor remains low until late in development, while in *H. erectus* this was already possible few days post birth, arguing for a high rigidity of the seahorse’s body armor in early stages compared to those of studied pipefishes. Still, in *S. typhle* we observed that some of the spaces between these bony plates become filled by much smaller, secondary plates without joints in adults, which do not arise from (or feature) previously described ridges (see also ^43^), arguing for the importance of a body armor that covers the body fully.

**Figure 14:**
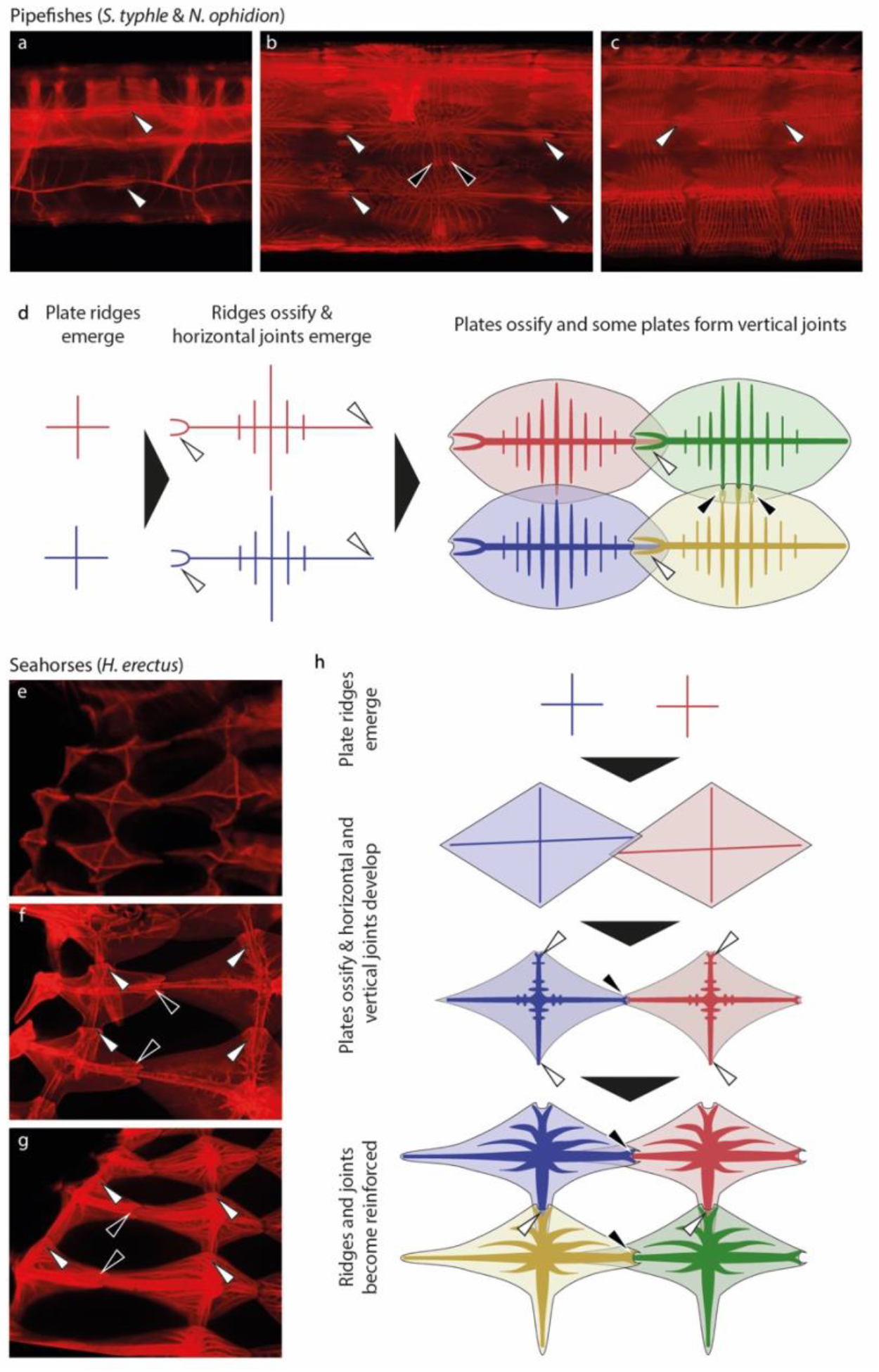
Post-natal development of the syngnathids’ bony plates. (a-c, e-g) Alizarin red fluorescence (stains ossified tissue) visualizes the development of bony plates, lateral view. Juvenile (a) and adult (b) *N. ophidion* (here: trunk region) and (c) adult *S. typhle* (here: caudal region; all cranial leftwards & ventral downwards. White arrow heads: longitudinal spine-and-groove joints; black arrow heads: smaller (or absent) transversal joints. *H. erectus* neck regions of an individual few days post birth (e), few weeks post birth (f) and adult (g; cranial to the top, ventral leftwards).

Finally, in *H. erectus* the cross-shaped condensations that form the bony plates are often elevated upon spine-like protrusions (primarily on the most dorsal plate row; not to be confused with spine-like protrusions found in *N. ophidion*), visible from ∼12dpm onwards (Fig. 10 lower panel, c,f). Interestingly, these plate protrusions emerged consistently among individuals first on the precaudal vertebrae three, five, seven and nine, while protrusions on the other precaudal vertebrae only emerged much later and remain smaller (hypertrophic vs. hypothropic plate protrusions, respectively). Caudally to the dorsal fin, such hypertrophic protrusions are found every three to four vertebrae, with individuals varying in patterns: some have hypertrophic protrusions consistently on every third caudal vertebra, some on every fourth, and others have more variable patterns. The difference between hypertrophic and regular spine-like protrusions translates also in differences in bony plate sizes, which are most notable in juveniles in *H. erectus*, while in adults the differences are much less pronounced (but still recognizable).

#### Pectoral fin & girdle morphology: loss and degeneration

The characterization of the morphology and developmental history of the syngnathids’ pectoral girdle and fin has been challenging due to their high degree of ossification as well as bone fusions and resorptions in juveniles and adults. Furthermore, the unusual positioning and shape of the endochondral disk derived bones in these fish led to some incorrect conclusions in previous studies (Fig. 15).^16^ The adult pectoral fins of *S. typhle* and *H. erectus* are highly derived and characterized by fusion with the bony plate armour and resorption of central fin parts. In *N. ophidion*’s rudimentary pectoral fins are only present in embryos and larvae, but are resorbed upon transition into juveniles (Fig. 11 c-g).

**Figure 15:**
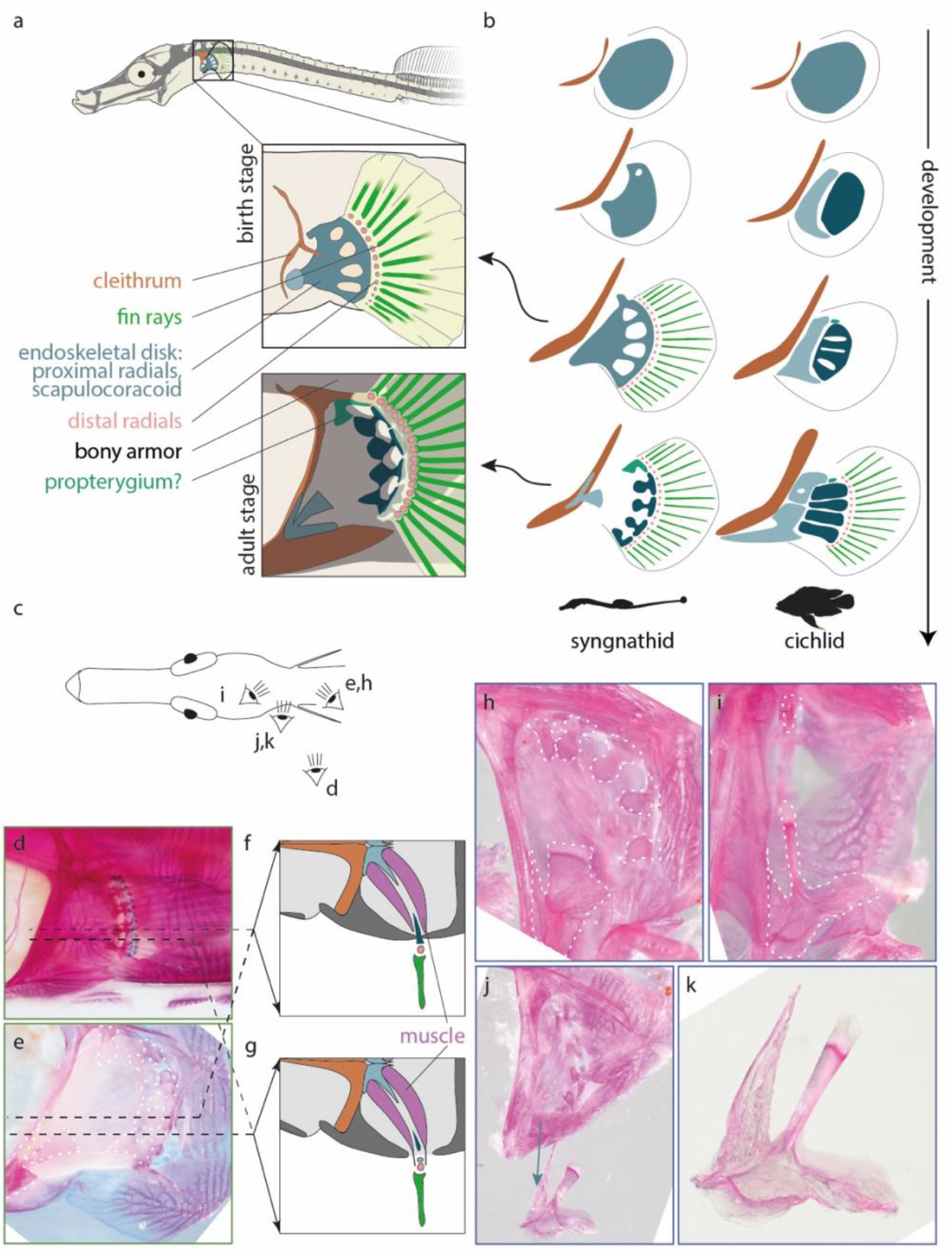
Pectoral fin base skeleton in syngnathids. (a) main skeletal structures contributing to pectoral girdle and fin development in *S. typhle*. (b) Comparison of main pectoral fin skeletal elements across developmental stages and between species; cichlid development following.^13^ (c) Viewing angles of plots d (left pectoral fin), e,h-k (right pectoral fin/girdle) after dissection. (d,e,h-k) Alizarin Red and Alcian Blue stains of adult *S. typhle* (d,e) or *H. erectus* (h-k). (f,g) Schematic dorsal view on the bone architecture of the pectoral fin at two different positions on the pectoral girdle (lateral downwards, medial upwards). Soft tissue attached to the scapulocoracoid and radials not removed (e) or removed (h-k). (h-k) Detailed view on the scapulocoracoid’s position and its morphology.

In all three species, pectoral fin bud condensation could be observed early on in development (Table 2) and an endoskeletal disk (giving rise to the scapula, coracoid, propterygium and proximal radials) could be recognized via Alcian Blue staining from 14, 15 and 6dpm in *N. ophidion*, *S. typhle* and *H. erectus*, respectively (Fig. 16 all panels, a). A main difference to most other teleosts is the failure of the endoskeletal disk cartilage to separate the initial cartilaginous Anlage into discrete scapula, coracoid and proximal radial elements during pre-release development. In *S. typhle* and *H. erectus*, a continuous Anlage emerges oriented almost perpendicularly to the body axis. Along its proximal ridge a forked thickening corresponds to a scapulocoracoid, which initially remains distally fused to the distal part of the endochondral disk (Fig. 15b; Fig. 16 mid and lower panels, e-h). In other teleosts, this more distal part of the endochondral disk gives rise to the proximal radials through cartilage resorption, however, in syngnathids this process appears to be partially suppressed. In *S. typhle* and *H. erectus* initial central fissures form, but these fail to extend proximally and distally before release. The recognizable five radial elements (the most dorsal one probably corresponding to the propterygium) remain fused distally and proximally connected to the scapulocoracoid – a conformation retained until bony plates covering the pectoral girdle have ossified sufficiently weeks to months after release (Fig. 15).

**Figure 16:**
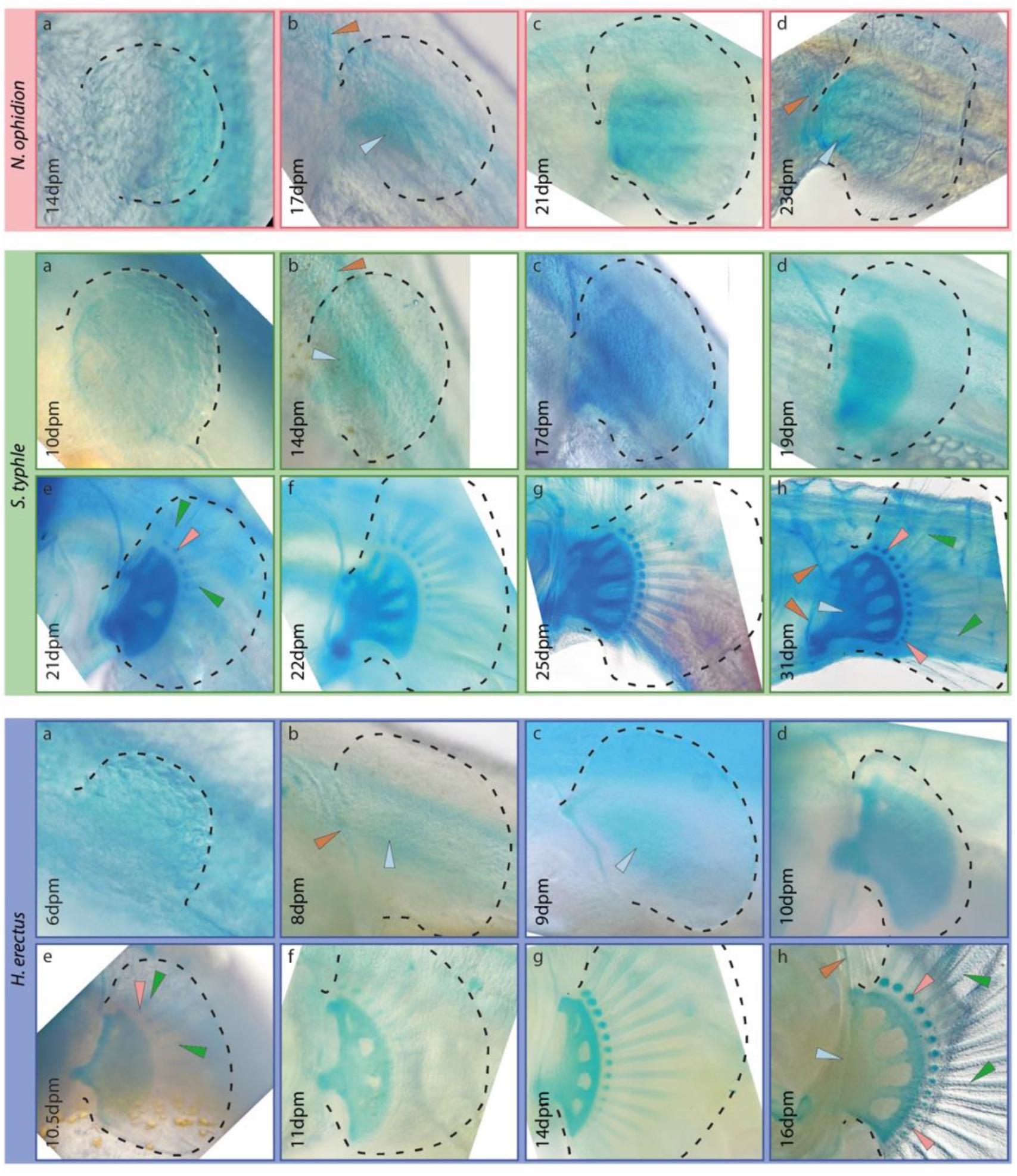
Pectoral fin chondrogenic development in examined syngnathids. Upper panel: pectoral fin development in *N. ophidion* is neotenic. Mid panel and lower panel: pectoral fin development in *S. typhle* and *H. erectus*, respectively, is similar across pre- and postnatal development; lower panel (h) is transversal view after surgical removal of trunk; dpm = days post mating.

Distal radials and fin rays appear as *de novo* condensations approximately 20dpm and 10dpm in *S. typhle* and *H. erectus*, respectively, and resemble in their ontogeny and structure those of other teleost fins^9^. Furthermore, we observe the formation of the cleithrum in a very similar manner. In adult *S. typhle* and *H. erectus*, the pectoral fin complex is attached to, and largely obscured by, strongly ossified tissue (Fig. 15 d) and its morphology is therefore difficult to observe. Dissections, reveal the presence of seemingly five proximal radial elements, which have now become attached to the dermal armor (Fig. 15 d-g). At their distal end proximal radials remain connected to each other via cartilaginous tissues and articulate with the fin rays (lepidotrichia) through the distal radials. Only a ventroproximal rudiment remains of the scapulocoracoid (Fig. 15 h-k) and proximal radials are individualized proximally. The radial elements do thus not directly articulate with the girdle but are fixed in place by their attachment to the bony armor, however, the two are connected through the dorso-ventral fin musculature that bridges the large intervening gap. The bony plates associated with the rudimentary girdle and radials probably provide the structural support that allowed the evolutionary reduction of radial and girdle elements. Thus, in adult *S. typhle* and *H.* erectus the pectoral fin/girdle is a highly derived complex wherein five radial-like elements are attached to the bony armor posterior-distally and the girdle is present as a small scapulacoracoid element anterior-proximally attached to the cleithrum and dermal plates. Interestingly the presence of five, instead of four, radials might equally be interpreted as a derived departure from the canonical Bauplan of the teleost fin.^45^ We, however, speculate that the most dorsal (rudimentary) radial element corresponds to the propterygium, whose formation and subsequent secondary fusion with the coracoid can also be observed in cichlids.^13^

Altogether, and while the homology of the various elements of the pectoral fin cannot yet be resolved with complete certainty, it appears that the autoapomorphic pectoral fin of *H. erectus* and *S. typhle* results from a failure to separate the initial endochondral disk into the separate components of the ancestral teleost pectoral fin, followed by a resorption of most of the initial cartilaginous *Anlage*.

An even more extreme developmental modification is found in *N. ophidion*, where pectoral fin development is truncated at the endochondral disk stage. During embryogenesis a single continuous endochondral disk Anlage connected to a fin fold is formed (Fig. 16 upper panel). However, this disk never shows any fissures nor do distal radials or fin rays develop. Instead, pectoral fin differentiation is terminated after approximately 21dpm (after which only the fin fold is still allometrically growing). Larval *N. ophidion* use these neotenic pectoral fins throughout their first post-release life stages (approx. the first two weeks post hatch), but lose them finally while transitioning into juveniles, which only possesses a dorsal fin after loss of the neotenic pectoral fins (Fig. 11 c-g).

#### Development of the caudal axial skeleton, dorsal fins and anal fins

The transition from pre-caudal to caudal vertebrae is indicated by the presence of the haemal arch in caudal vertebrae, which forms from protrusions found at 18 and 9.5 dpm in *S. typhle* and *H. erectus*, respectively, while these are not present yet in *N. ophidion* at hatching (Fig. 17; at 10 days post hatch they could be observed using Alcian Blue staining). The dorsal fin develops from mesenchymal condensations, which form the proximal radials (pterygiophores), after which distal radials and rays form. In *N. ophidion*, dorsal fin differentiation is not concluded before hatching, likely because the species relies on its well-developed fin fold for locomotion during the first days post hatching, but at approximately 12 days post hatch rays were completely formed. Seahorses rely for their locomotion on dorsal fin movement.^46^ In line with this central importance seahorses complete dorsal fin formation relatively earlier in development, even compared to *S. typhle*. In *S. typhle* and *H. erectus* the anal fin forms fairly synchronized with the dorsal fin (Table 2).

**Figure 17:**
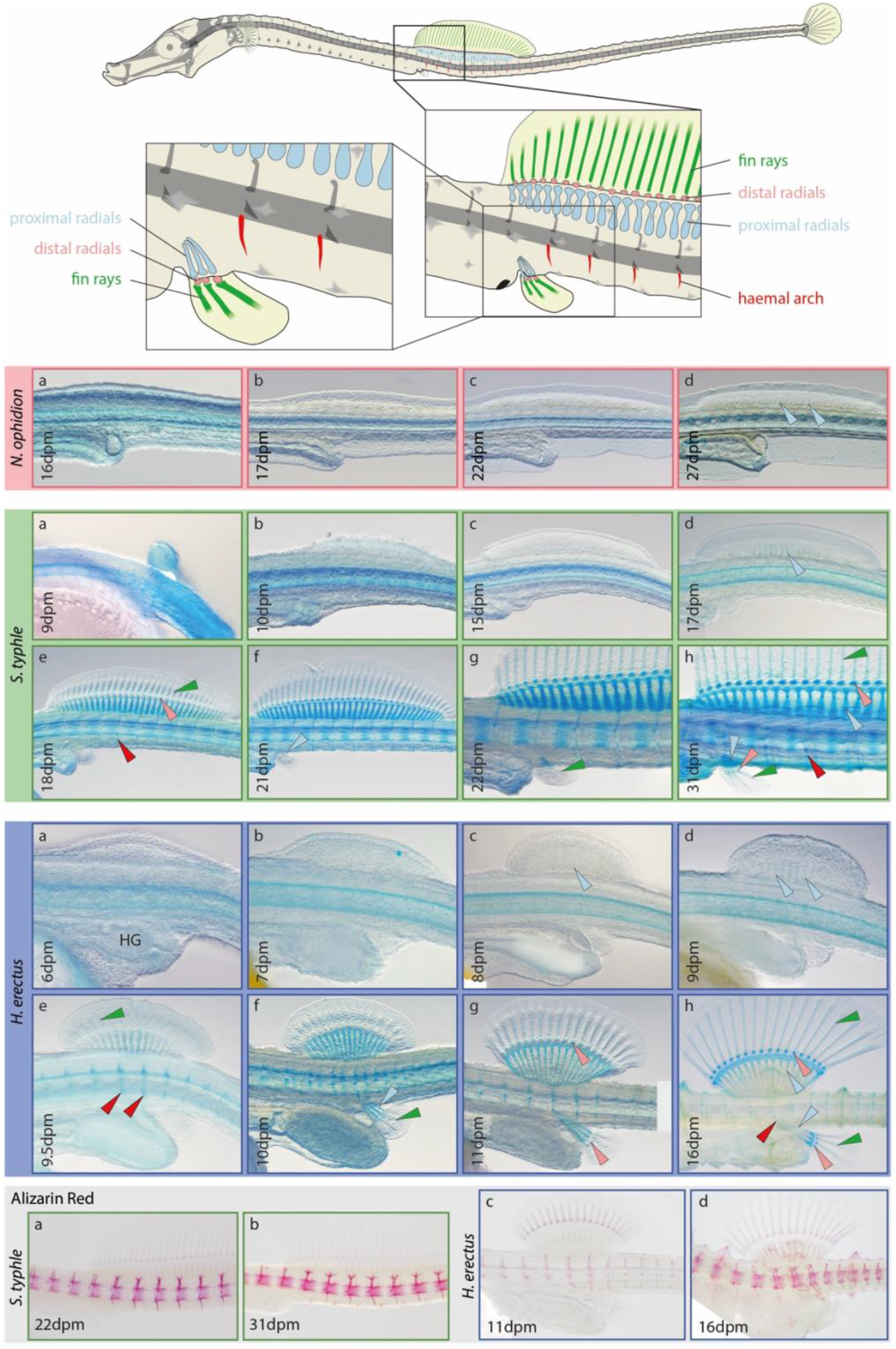
Dorsal, anal fin and haemal arch skeletal development in examined syngnathids. Upper panel: main skeletal structures contributing to dorsal and anal fin development in *S. typhle*. Second panel: dosal fin development in *N. ophidion* is not concluded by hatching day. Third panel and forth panel: dorsal and anal fin development are almost synchronized in *S. typhle* and *H. erectus*, respectively, and similar between species. Lower panel: ossification patterns in *S. typhle* and *H. erectus* (*N. ophidion* did not show any staining in this region before hatching); dpm = days post mating.

#### Ural region and the caudal fin

The teleosts’ caudal fin is a complex structure and, in contrast to the other fins, its endochondral components are derived from the vertebral column. In the caudal fin, rays articulate directly with highly modified neural and haemal arches (the “epurals” and “hypurals”, respectively), which originate from the most posterior vertebrae (for overview see ^13^). In syngnathids, the caudal fin was lost in several taxa (often in favor of a prehensile tail),^42^ and in those with caudal fins it appears highly derived (Fig. 18). From among the many individual bony structures found in most other teleosts, only the urostyle (or a urostyle-like most posterior element) could be identified with some certainty in *S. typhle* and *H. erectus*, although in the latter it does not retain its initial dorsal bending (visible at 8dpm), which would be typical for teleost fishes. In *S. typhle* the formation of a single hypural cartilage can be observed (18dpm). Its partially bifurcated shape is suggestive of specification into a dorsal and a ventral domain. This cartilage does however not separate into individual hypural radials. Actinotrichia can already be observed at 17pdm and forming fin rays directly articulate with the hypural cartilage (starting approx. 19dpm). This morphology is still present in adult *S. typhle*, however, it should be noted that bony plates cover the entire tail up to the caudal peduncle, presumably providing structural stability to the caudal complex (similar to the pectoral fins). In *S. typhle* the presence of an additional transient dorsal Anlage can be detected during development (15-19dpm), which possibly corresponds to a rudimentary uroneural (Fig. 18). In *H. erectus*, the caudal fin is degenerated (as reported before for other *Hippocampus* ^16,18,47^) and although some hypural cartilage is formed, it does not attain a shape resembling the caudal fin in *S. typhle* (Fig. 18 fourth panel). The outgrowth of 2-5 developing rays, as suggested by actinotrichia, stagnates soon after its onset and these are not recognizable anymore in adult *H. erectus* (see also *in situ* hybridization section below).

**Figure 18:**
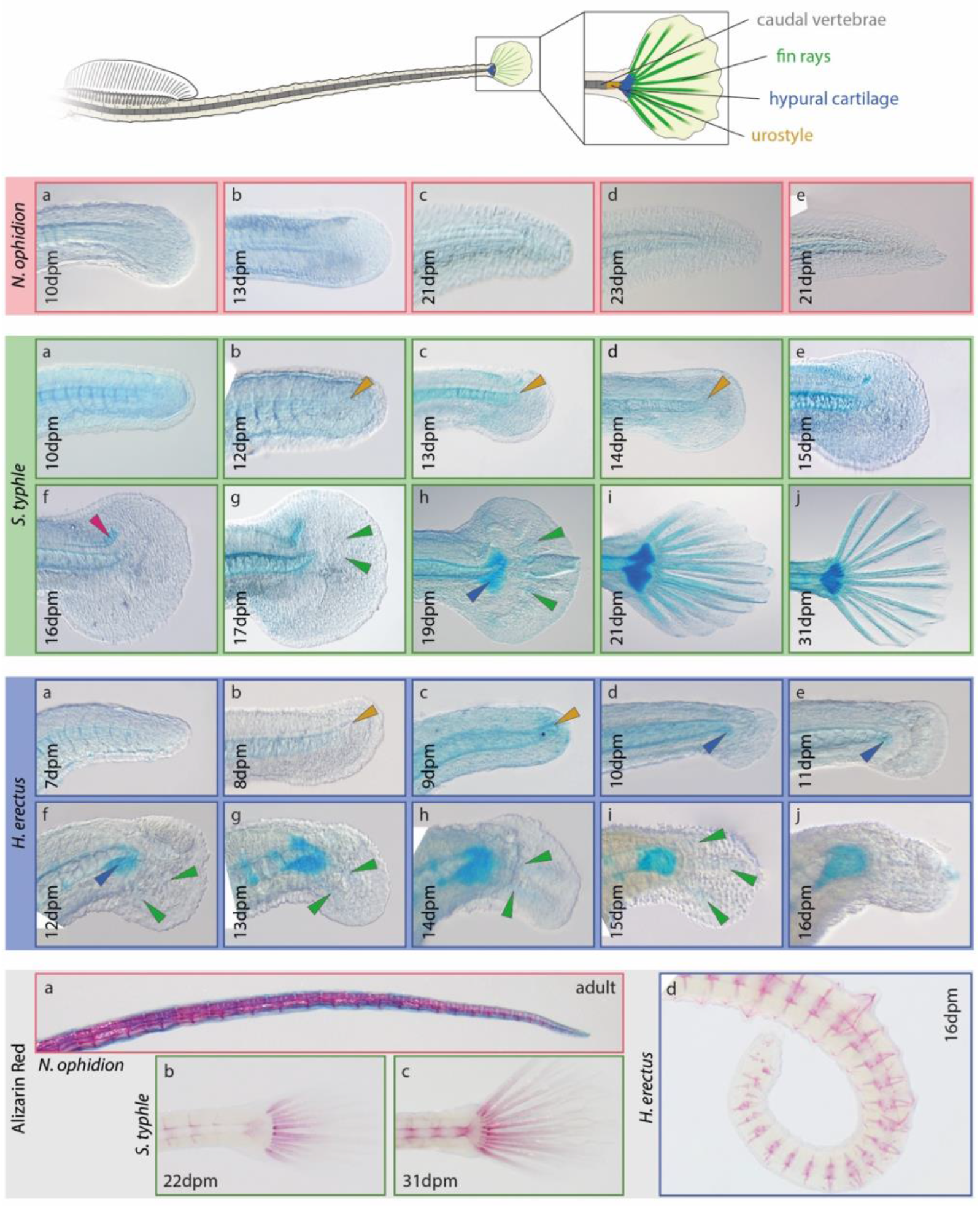
Ural region pre-natal skeletal development in examined Syngnathids. Upper panel: main cartilages contributing to caudal fin development in *S. typhle*. Second panel: *N. ophidion* develops no caudal fin but instead a pronounced fin fold, which weeks post hatch is resorbed and the tail becomes prehensile. Third panel: caudal fin development in *S. typhle*. The red arrow marks potentially a rudimentary uroneural. Forth panel: caudal fi development in *H. erectus* is not concluded and only a rudimentary caudal fin is formed on the prehensile tail. Lower panel: ossification in the ural region; dpm = days post mating, second to forth panel Alcian Blue staining; lowest panel Alizarin Red staining (and Alcian Blue in (a)).

Ossification in the most posterior spine region of *N. ophidion* was only detected in subadult fish, with fairly uniform vertebrae merely decreasing in size forming the caudal tip (Fig. 18 lower panel, a). In contrast, ossification of the hypural cartilage commences at 22dpm in *S. typhle* with rays starting to ossify already one to two days earlier (Fig. 18 lower panel, b). In *H. erectus*, the hypural cartilage did not undergo ossification before release, however, alizarin red stains show how the very organized patterns of one neural and haemal spine per vertebra typical for the rest of the caudal region is disrupted in the most posterior vertebrae, illustrating the overall disarray of regulatory processes orchestrating the ural region’s development in this species.

#### Col10a1a reveals particularly active zones of cartilage and bone formation

Teleost *col10a1a* is expressed in differentiating chondrocytes and osteoblasts and thus *in situ* hybridization can reveal the developmental onset of such cartilaginous and bony skeletal structures. In the examined stages, *col10a1a* expression was first detected in the cleithrum, revealing the presence of this structure already after 12 and 7dpm in *S. typhle* and *H. erectus*, which is considerably earlier than Alcian Blue staining suggested (Fig. 19). Subsequently, *col10a1a* expression was detected in Meckel’s cartilage, the hyosymplectic, retroarticulare and the frontale (after approx. 16 and 10dpm in *S. typhle* and *H. erectus*, respectively). After these stages, *col10a1a* expression was primarily detected in the outgrowing fin lepidotrichia (even in the rudimentary caudal fin of *H. erectus*; Fig. 19 lower panel, f-h) and also in the center of emerging bony plates in *H. erectus* 14dpm.

**Figure 19:**
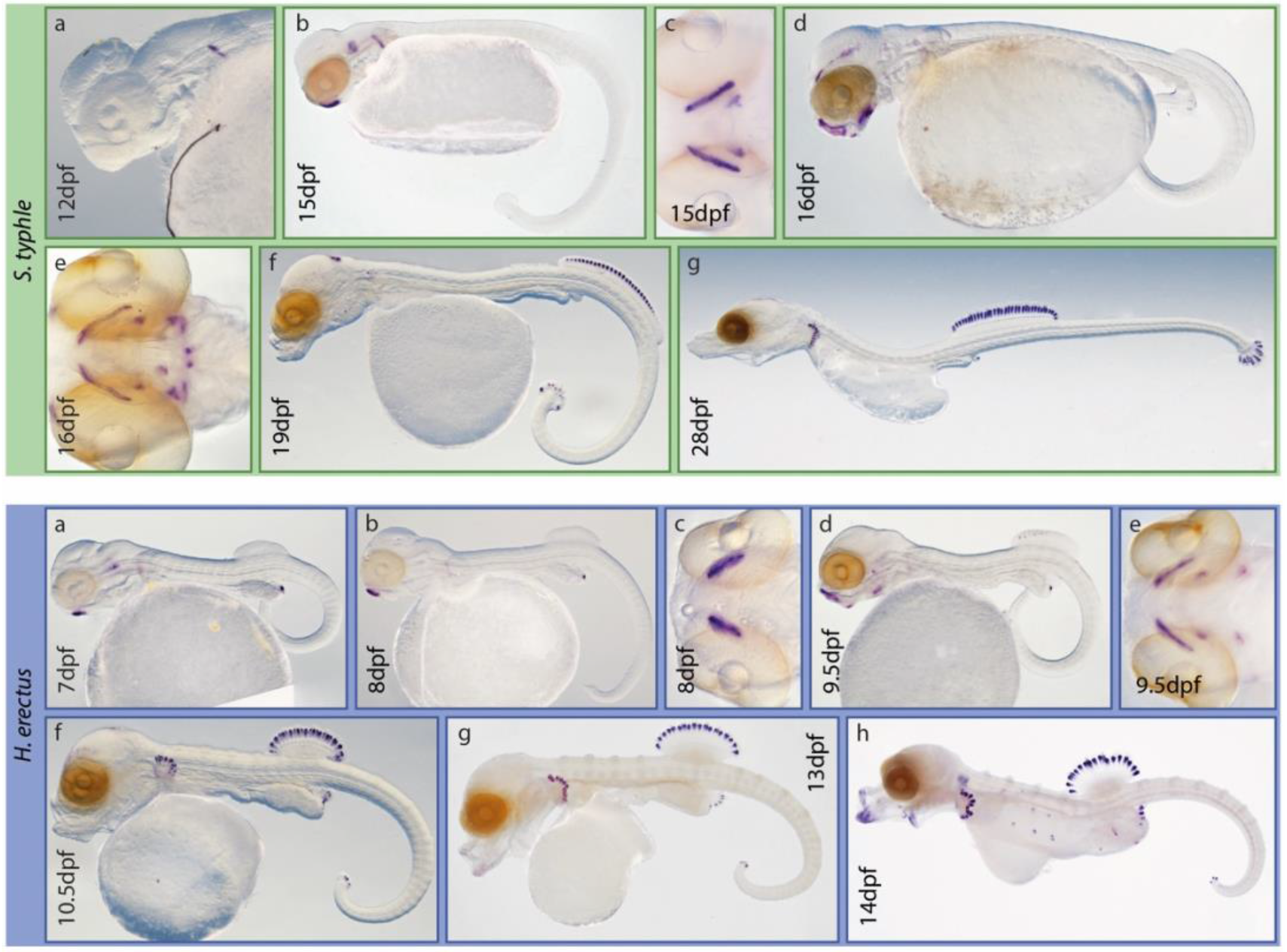
Skeletogenesis in two examined syngnathids as revealed via *col10a1a* in situ hybridization. Upper and lower panel: onset of cartilage/bone formation in *S. typhle* and *H. erectus*, respectively. All lateral view, except (c) and (e), which are ventral view. Expression of c*ol10a1a* suggests chondro-skeletogenesis commences in the pectoral girdle, followed by jaws and fin rays. Interestingly, also bony plate precursors show some expression; dpm = days post mating.

### Myogenesis in syngnathids as revealed by *in situ* hybridization

Seahorse skeletal musculature has been investigated in conjunction with the fish’s unique grasping tail in previous studies.^42,48^ We investigated the early development of such muscle using *in situ* hybridization for *mybpc1*, a gene expressed in slow striated skeletal muscle ^49^ that proved to be a useful marker gene for all skeletal muscle in this study (see also Experimental Procedures; Fig. 20).

**Figure 20:**
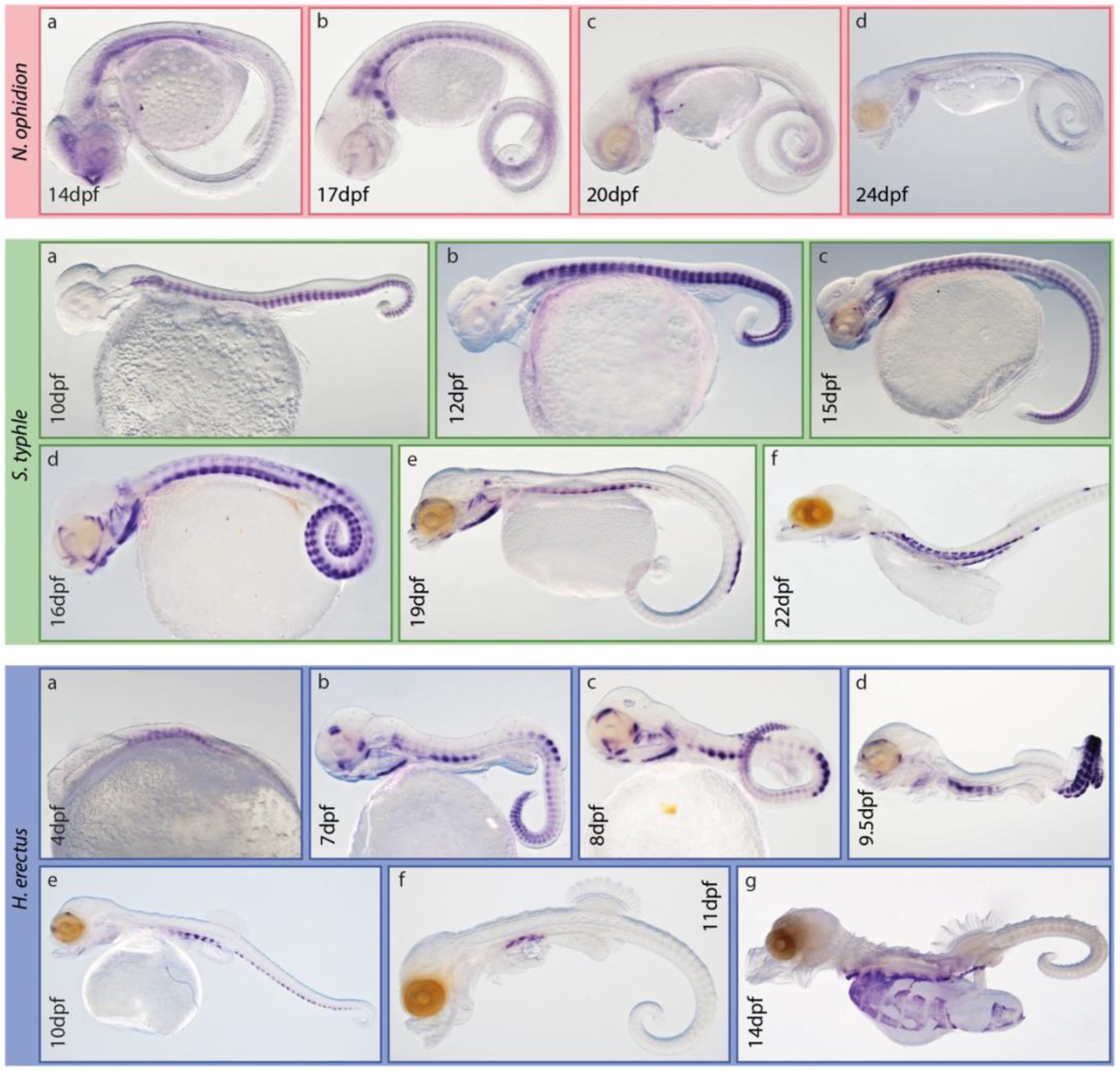
Formation of slow skeletal muscle in examined syngnathids as revealed via *mybpc1 in situ* hybridization. Upper, mid and lower panel: skeletal muscle formation in *N. ophidion*, *S. typhle* and *H. erectus*, respectively. *Mybpc1* expression suggests that myogenesis commences with axial musculature and epaxial and hypaxial expression patterns vary between species. Head musculature follows and appears to be concluded before release. All lateral view; dpm = days post mating. Lower panel at 4dpf without head.

Patterns of skeletal myogenesis are similar across examined species and commences with the axial musculature and myomeres form adjacent to the somites (present at 10dpm and potentially before in *S. typhle*; Fig. 18 mid panel, a). Subsequently, pronounced *mybpc1* expression outlines the sternohyoideus muscle, as well as the onset of myogenesis of branchial, extraocular, neck-region and jaw musculature (between 12 and 15dpm in *S. typhle*). At this stage expression patterns of *mybpc1* also suggest that axial musculature is separated into the hypaxial and epaxial portion, which will be separated by the transversal process upon its formation. Gene expression suggests continuous head musculature formation throughout the next few days (at 20dpm head *mybpc1* expression in the head is virtually absent in *S. typhle*, except in the sternohyoideus muscle). During this period, expression patterns in the axial musculature become more differentiated, suggesting an earlier conclusion of myogenesis of epaxial compared to hypaxial musculature in the trunk (with the exception of the myomeres directly anterior to the anterior border of the dorsal fin, where *mybpc1* expression remains strong). In the caudal region, *mybpc1* expression attenuates first anteriorly, with posterior myomeres retaining higher expression levels, which likely reflects their later onset of development. While in *S. typhle mybpc1* expression attenuates first in the hypaxial caudal musculature, *H. erectus* shows the opposite pattern, with the epaxial musculature first concluding its development. In late embryonic stages *mybpc1* expression is only detected in the hypaxial trunk musculature that is likely involved in the closure of the body wall once the yolk sack is sufficiently resorbed. When larvae/juveniles are released, distinct *mybpc1* expression could not be detected anymore (data not shown), arguing for a conclusion of myogenesis onset prior to this event. In *N. ophidion*, *mybpc1* expression patterns were less distinct but available data suggest a similar myogenesis pattern to *S. typhle*, but again with a substantial delay in relative pre-natal timing, as, for instance, the *mybpc1* expression patterns of *N. ophidion* at 24dpm (approx. 89% of pre-natal development concluded) corresponds to a stage found at 22dpm in *S. typhle* (approx. 71% of pre-release development concluded).

## Discussion

Syngnathids are extremely derived fishes featuring some of the most bizarre phenotypes found in teleosts, such as their fin loss, very unusual body shapes, paternal care and highly modified immune systems.^6–8^ Many of these iconic traits develop before the embryos hatch (or are born) and thus only few studies have so far attempted to describe the developmental trajectory turning a seemingly generic fish embryo into an evolutionary extravaganza.

### Comparative development of studied syngnathids: direct vs. indirect development

Studied syngnathids’ exhibit quite distinct adult morphologies: *N. ophidion* being extremely elongated with a small head and almost no fins, *S. typhle* with a more generic pipefish body morphology, and *H. erectus* with the upright posture of the seahorses. During development, however, the smaller evolutionary distance and possibly the more similar mode of parental care between *S. typhle* and *H. erectus* becomes apparent,^26,50^ as their development is more synchronized and more pronounced differences between the two appear relatively late in development (Table 2). Both species also have a rather direct development, with juveniles released during birth relatively closely resembling their adult forms, and all characteristic body features (such as the adult set of fins and approximate body proportions) have been (or are being) developed. This stands in stark contrast to *N. ophidion*, which appears to hatch in a less developed state compared to *S. typhle*, that has a similarly long pre-natal period (Table 1). Furthermore, freshly hatched *N. ophidion* do not feature the characteristic adult traits: the head is still relatively large with a long snout, pectoral fins are still present and a well-developed fin fold is used for locomotion. Instead, adult characteristics are slowly developed approximately 2-6 weeks after hatching. Therefore, *N. ophidion* presents itself as an indirect developing species with a pronounced larval stage, while the development of *S. typhle* and *H. erectus* is much more direct, possibly as a result or by-product of their more derived form of male pregnancy.^5,6,13^ Thus, developmental stage upon release may be linked to the lineage (with *Nerophinae* featuring a larval stage and *Syngnathinae* not), pregnancy type (with *Nerophis* likely not providing nutrients to embryos, while this is more likely/shown for (some) Syngnathinae^51,52^), or other factors not considered here and /or their interactions.

### Development of characteristic syngnathid traits

The snout-like jaws of syngnathids are among their most characteristic traits. We find that the onset of snout elongation is primarily driven by the elongation of the hyosymplectic cartilages and the unfolding of the ethmoid cartilage (which arises in a curved configuration) during the second half of pre-natal development, and thus prior to the onset of ossification of these elements. By rotation of the palatoquadrate to an almost perpendicular position to the hyosymplectic the tube-like buccal cavity is formed (Fig. 10), which crucially facilitates pivot feeding.

Bony dermal plates are another characteristic of syngnathids that is also found in other fish families, such as sticklebacks, and while the homology of plates across these clades was never established, their occurrence contributed to their (no longer accepted) grouping into the Gasterosteiformes.^53^ Here, we find that syngnathid bony plates arise from cross-like tissue condensations late in pre-natal development (Fig. 12-14). As we observed that – within each plate row – per vertebra exactly one bony plate is formed, we infer that their ontogeny is closely linked to the segmented structure trunk as provided by the somites.^13^ Yet, the exact homology of bony plates across fishes, and/or whether evolutionarily available gene networks were co-opted during their evolution, remains unknown. Furthermore, it remains unclear what determines the number of plate rows along the trunk and tail region of the fish. Finally, as plates show a variety of different joint types, exploring the mechanisms determining which joint type forms during development, and understanding if these joint types are genetically predetermined or if joint development is (partially) informed by mechanical interactions among neighboring plates during development, would further our understanding of how complex novel morphological structures can evolve.

Among syngnathids, the presence or absence of fins appears to be a particularly diverse characteristic and for any single fin that is found among syngnathids (e.g., the fin set that *S. typhle* has) there are representatives that have lost that particular fin (or multiple). For instance, in *N. ophidion*, we did not identify any evidence of a (rudimentary) caudal fin being formed even early on in development, suggesting that the regulatory molecular network underlying caudal fin formation is dysfunctional at an upstream position, leading to a complete fin loss (similar to pelvic fin loss across all syngnathids ^8^; Fig. 18). In contrast, in *H. erectus*, we (as many before, e.g., ^16,18,47^) have observed the onset of fin development while generic fin differentiation and outgrowth failed, arguing for a downstream dysfunction that led to a functional fin loss.

Another example of fin loss could again be observed in the indirectly developing *N. ophidion*, where larval pectoral fins are lost during the metamorphosis several weeks after hatching. In these fish, pectoral fins and a well-developed median fin fold along the tail are resorbed (and also the head shape changes, e.g., the snout becomes shorter; Fig. 11 & 15). A similar, but even more extreme metamorphosis has been described before in *Bulbonaricus brucei* (pug-nosed pipefish),^54^ in which even the larval dorsal fin is resorbed (and facial alterations are more extreme, leading to a pug-like facial appearance). Our study reveals that in *N. ophidion*’s generic pectoral fin development was never actually concluded, despite the pectoral fins are used by the larvae during their first weeks post hatching. Instead, pectoral fin development was seemingly truncated before skeletal differentiation occurred, and while fin outgrowth in size continued, the fin morphology clearly remained neotenic (Fig. 16 upper panel). Similar to the caudal fin loss in *H. erectus*, this argues for a rather downstream dysfunction in the generic pectoral fin regulatory network. Such pattern of aborted development resembles digit loss in cows or birds where the Anlagen of the missing digit are specified during embryogenesis but fail to develop into a lasting adult structure.^55,56^ Exploring evolutionary shifts in musculoskeletal allometric growth patterns (e.g. via heterochrony) might thus be a fruitful approach to understand the evolutionary trajectories underlying some of the most iconic syngnathid traits, such as the prehensile tail or craniofacial differences (as suggested in ^18^). Finally, another interesting avenue for further research is to understand how metamorphosis induces the substantial developmental remodeling observed in maturing *N. ophidion* (and more so in pug-nose pipefish) and identify (hormonal) triggers, such as thyroid hormone,^57^ leading and controlling this process.

### Are seahorses developmentally truncated pipefish?

An important derived feature of seahorses is their characteristic upright swimming position, whereby their trunk is oriented vertically, while the head is oriented not upright but horizontally (or even more bend towards the trunk; Fig. 1 d). This bend posture, whereby the snout is at a sharp angle to the trunk can be observed early during ontogeny, around. 11 dpm (Fig. 3-5). Although the swimming posture of pipefishes is typically horizontal and their head and trunk are aligned, they pass through a seahorse-like developmental stage, e.g., *S typhle* 21 dpm, during which head and trunk form a 90-degree angle. This would suggest homologous bodyplan postures during development but a failure of seahorses to complete the process of straightening their head-trunk connection, i.e., developmental truncation of an ancestral ontogenetic trajectory. As discussed before, another potential indication for developmental truncation during seahorse development as compared to their sister pipefish clade (here: *S. typhle*) is the abortion of caudal fin development. In the latter case, it is clear that an Anlage forms which proceeds till the stage of fin ray condensations (as shown by *col10a1a in situ* hybridization). The outgrowth is, however, precociously terminated leading to the absence of caudal fins in adults. Therefore, our ontogenetic comparison between *H. erectus* and *S. typhle* development provides several indications that the unique body plan of seahorses, could in part result from developmental truncation.

### Concluding remarks

Syngnathids feature some of the most bizarre traits found among fish, such as bony plates, snout-like facial anatomy, male pregnancy, highly modified immune systems and many others.^5,7,8,42^ For some of these traits, understanding the developmental trajectories syngnathids undergo is vital to unravel the evolutionary mechanisms and ecological & life-history events affecting trait development. Our study provided such insights and despite of being focused on only the early-life development of few selected syngnathids, we recognized in our data a plethora of potential evolutionary mechanisms shaping specific aspects of syngnathid evolution and development, and believe that particularly heterochrony may be underlying some of the morphological variability found in this extraordinarily diverse family of fish.

## Experimental Procedures

Fish used in this study were kept at the fish keeping facilities of the Helmholtz-Centre for Ocean Research Kiel (GEOMAR). Pipefishes were originally collected in a shallow, close-by eelgrass meadow; coordinates: 54.391103, 10.191419; seahorses were obtained from commercial fish breeders, and sampling occurred from March to August 2020. *Nerophis ophidion* and *Syngnathus typhle* stock individuals were kept in community tanks at 16-18°C and *H. erectus* at 23°C (all +/-0.5°C) and inspected daily for pregnancies. When a pregnancy was detected, for *N. ophidion* and *S. typhle* the individual was separated and kept at constant 16 °C in a smaller tank connected to a water circulation system, while *H. erectus* were kept in their stock tanks as individuals could easily be identified. *S. typhle* and *H. erectus* were fed live and dead mysids, while *N. ophidion* received live *Artemia* nauplii and live mysids, and all food was enriched with vitamins and fatty acids. Measurements taken in this study reflect individuals kept and raised at these temperatures and developmental times may differ when fish are raised at other temperatures (temperature affects developmental speed ^58,59^).

For sampling, pregnant *S. typhle* and *H. erectus* individuals were killed using MS222 and subsequently decapitated. The brooding flaps/pouch were opened and eggs/larvae were removed. For *N. ophidion*, eggs could easily be removed from the fathers’ bellies using brushes, and males could be sampled several times in this manner, providing a continuous series of eggs with a known age difference. Obtained eggs and larvae were either immediately killed using MS222, photographed and fixed (using 4% PFA in PBS, overnight, and then gradually moved to MeOH; some photos were taken post fixation and MeOH storage) or, in the case of selected *H. erectus* embryos, cultivated *in vitro* in petri dishes at the same temperature as their parents until sampling. Note that the size of the yolk sac prior to full gastrulation was affected by the osmolarity of the water in which the eggs were submerged for photography (filtered Baltic sea water of ∼17ppm). For culturing embryos, filtered North Sea seawater diluted to a salinity of ∼20ppm was used, as embryos (age 1-7 days) showed highest average survival at this salinity. For embryos older than 12 days, salinity was gradually increased to meet 35ppm after 16 days. As mating could only be observed for few *H. erectus* individuals during this experiment, intermediate stages were obtained by rearing offspring *in vitro*, as described, and daily sampled. In *H. erectus*, while stages for each day in development were identified, they likely not represent 24h intervals each time (e.g., day nine to ten, and ten to eleven are likely >24h), possibly due to slightly slower growth rates of embryos *in vitro* and because the onset of embryonic development was never exactly known, as fertilization occurs inside the pouch. Therefore, the manuscript uses the unconventional “hours/days post mating” to indicate the age of embryos. While in many teleosts the time of mating and fertilization coincides, in syngnathids this is likely not the case and fertilization happens some time (potentially hours) after mating.

Skeletal stains were performed on fixed embryos/larvae using the following staining solution (for 1 ml of “low-acid double-stain”): 12.5µl Alcian Blue solution (0.4% in 70%EtOH), 50µl acetic acid (100%), 57.5µl MgCl_2_ (2M), 870µl EtOH (70%), 10µl Alizarin Red solution (0.5% in H_2_O), based on ^40^. Stains were performed typically for three to four hours, but larger individuals (i.e., free-swimming larvae, juveniles, adults) were kept for up to 12h in staining solution. Treating embryos for even 15-20 minutes in the low-acid staining solution dissolved all subtly ossified structures and led to a delayed detection of mineralization (very labile mineralizing was already noted by ^24^ for syngnathids), and additional Alizarin Red staining did not recover these patterns, but only led to strongly stained yolk sacs (see Fig. 5 mid panel, g). Also, as acid-free double staining solution (according to ^40^) did not yield skeletal double-stains of sufficient quality even after optimization, staining for Alizarin red was performed (10µl Alizarin Red solution (0.5% Alizarin Red in H_2_O) per millilitre 70% EtOH) on separate individuals. After staining, individuals were repeatedly washed in 70% EtOH, gradually transferred to H_2_O, and then bleached and cleared using 0.5-1.5% H_2_O_2_ in 0.5-1% KOH, depending on pigmentation stage for 5-30 min; juveniles and adults longer, and continued to be clear in 0.5% KOH in 50% glycerol for several hours to days.

Photographs of embryos were taken using a Nikon SMZ18 stereomicroscope system equipped with a DS-Fi3 camera. For positioning purposes, samples were typically photographed on agarose gels with indentations for yolk sacs or fins, facilitating the proper alignment of the embryos. In early stages, eggs were de-chorionated whenever possible without damaging the embryo. Obtained photos were processed in Adobe Photoshop. The following parameters (or a subset of them) were adjusted in obtained images: brightness (“exposure”) and contrast, white balance, rotation and reflection, cropping, shadows/highlights (strongly pigmented eyes appear brighter), saturation, agarose-gel background noise removal and extension. For photos taken of fluorescent Alizarin Red, background fluorescence was removed by adjusting the color curve and overlays were also produced in Photoshop. All adjustments (except for the background) were always applied to the whole image and none was made with the intention to hide/distort any information of the obtained photos. For all images, raw file copies are available upon request.

Morphological structures were identified according to previously published work.^10,16,18,19^ The branchial basket could generally not be visualized well across all stages in this study and thus was not described, as it was done before in detail.^17,60,61^ Measurements (e.g., ray counts, length measurements) were only recorded in one or few individuals per time point, as many samples come from fish stocks that have been kept in the lab for generations and the genetic variability of these stocks is expected to be reduced compared to natural populations.

44 landmarks identified in ImageJ (v. 1.5.1) on lateral photos of heads were used to describe the shape of syngnathid heads and its key cartilages across developmental stages (photos across species and stages were randomized for this process; *N. ophidion* = 29, *S. typhle* = 42; *H. erectus* = 49; Fig. 10 a; Supplementary Table S1). As structures were not always identifiable/unambiguous at all stages, landmark locations were estimated when structures were not visible (i.e., not yet developed). This was necessary as the principal component analysis (PCA), which was employed for data analysis, requires a complete set of landmarks across all samples. Furthermore, several landmarks were put along a smooth structure contour (e.g., to describe the ethmoid cartilage bending, Fig. 10 a) and thus cannot be considered taxic homolog, but instead they were roughly put with equal distance to one another on the respective structure. This approach led in our evaluation to more accurate representation of shape homology when compared to treating the landmarks as sliding (semi-) landmarks. Due to these limitations, the geometric morphometric dataset was interpreted cautiously. A generalized Procrustes analysis was performed on obtained data-points (gpagen() function from “geomorph” package v. 4.1.2 ^62^) and a PCA was conducted in R (v. 4.1.3).^63^ A scree-plot suggested that most variation is loaded on PC1-3 (PC4 explains ∼2.3% of variation and is not considered here anymore).

*In situ* hybridization was performed to visualize the expression of *col10a1a* and *mybpc1* to monitor cartilage/bone and skeletal muscle formation, respectively.^64,65^ *Col10a1a* is in teleosts expressed both in differentiating chondrocytes and osteoblasts expression expressed,^64^ and facilitates the detection of cartilage and bone formation much earlier than staining techniques used in the manuscript. Thus, the two approaches complement each other. *Mybpc1* was described to be expressed in slow striated skeletal muscle ^66^ but proved in this study to be a useful marker gene using *in situ* analysis for overall early skeletal muscle development as it revealed distinct and unambiguous patterns. As gene trees confirmed that indeed *mybpc1* (and not its paralogs) was cloned to synthesize *in situ* probes, observed expression patterns suggest that *mybpc1* is also expressed in fast striated muscle development in syngnathids.

## Acknowledgements

We would like to thank Fabian Wendt for raising *N. ophidion* larvae, Beke Hansen for preliminary work on the geometric morphometrics data-set and Katharina Krüger for helping with skeletal stainings and their photography. This project was supported by the European Research Council (ERC) under the European Union ’s Horizon research and innovation program (MALEPREG: eu-repo/grantAgreement/EC/H2020/755659) to OR.

## Author Contributions

R.F.S., O.R., J.M.W and D.A. conceived the project. R.F.S. processed and analyzed samples and drafted the manuscript. All authors contributed via discussions in regular meetings, revised and approved the final version of the manuscript.

## References

1. Small C, Bassham S, Catchen J, et al. The genome of the Gulf pipefish enables understanding of evolutionary innovations. Genome biology. 2016;17(1):258.

2. Small CM, Healey HM, Currey MC, et al. Leafy and weedy seadragon genomes connect genic and repetitive DNA features to the extravagant biology of syngnathid fishes. Proceedings of the National Academy of Sciences. 2022;119(26):e2119602119.

3. Qu M, Liu Y, Zhang Y, et al. Seadragon genome analysis provides insights into its phenotype and sex determination locus. Science Advances. 2021;7(34):eabg5196.

4. Teske PR, Beheregaray LB. Evolution of seahorses’ upright posture was linked to Oligocene expansion of seagrass habitats. Biology letters. 2009;5(4):521–523.

5. Stölting KN, Wilson AB. Male pregnancy in seahorses and pipefish: beyond the mammalian model. BioEssays. 2007;29(9):884–896.

6. Wilson AB, Vincent A, Ahnesjö I, Meyer A. Male pregnancy in seahorses and pipefishes (family Syngnathidae): rapid diversification of paternal brood pouch morphology inferred from a molecular phylogeny. Journal of Heredity. 2001;92(2):159–166.

7. Roth O, Solbakken MH, Tørresen OK, et al. Evolution of male pregnancy associated with remodelling of canonical vertebrate immunity in seahorses and pipefishes. Proceedings of the National Academy of Sciences of the United States. 2020.

8. Lin Q, Fan S, Zhang Y, et al. The seahorse genome and the evolution of its specialized morphology. Nature. 2016;540(7633):395–399.

9. Van Wassenbergh S, Roos G, Genbrugge A, et al. Suction is kid’s play: extremely fast suction in newborn seahorses. Biology Letters. 2009;5(2):200–203.

10. Leysen H, Jouk P, Brunain M, Christiaens J, Adriaens D. Cranial architecture of tube- snouted gasterosteiformes (Syngnathus rostellatus and Hippocampus capensis). Journal of Morphology. 2010;271(3):255–270.

11. Praet T, Adriaens D, Cauter SV, Masschaele B, Beule MD, Verhegghe B. Inspiration from nature: dynamic modelling of the musculoskeletal structure of the seahorse tail. International journal for numerical methods in biomedical engineering. 2012;28(10):1028–1042.

12. Porter MM, Novitskaya E, Castro-Ceseña AB, Meyers MA, McKittrick J. Highly deformable bones: unusual deformation mechanisms of seahorse armor. Acta biomaterialia. 2013;9(6):6763–6770.

13. Woltering JM, Holzem M, Schneider RF, Nanos V, Meyer A. The skeletal ontogeny of Astatotilapia burtoni–a direct-developing model system for the evolution and development of the teleost body plan. BMC developmental biology. 2018;18(1):1–23.

14. Flegler-Balon C. Direct and indirect development in fishes—examples of alternative life-history styles. Alternative life-history styles of animals. Springer; 1989:71–100.

15. Balon EK. Alternative ways to become a juvenile or a definitive phenotype (and on some persisting linguistic offenses). Environmental Biology of Fishes. 1999;56(1):17–38.

16. Novelli B, Otero-Ferrer F, Socorro J, Caballero M, Segade-Botella A, Domínguez LM. Development of short-snouted seahorse (Hippocampus hippocampus, L. 1758): osteological and morphological aspects. Fish physiology and biochemistry. 2017;43(3):833–848.

17. Novelli B, Socorro J, Caballero M, Otero-Ferrer F, Segade-Botella A, Molina Domínguez L. Development of seahorse (Hippocampus reidi, Ginsburg 1933): histological and histochemical study. Fish Physiology and Biochemistry. 2015;41(5):1233–1251.

18. Franz-Odendaal TA, Adriaens D. Comparative developmental osteology of the seahorse skeleton reveals heterochrony amongst Hippocampus sp. and progressive caudal fin loss. EvoDevo. 2014;5(1):1–12.

19. Kadam K. The development of the chondrocranium in the seahorse, Hippocampus [Lophobranchii]. Zoological Journal of the Linnean Society. 1958;43(293):557–573.

20. Sommer S, Whittington CM, Wilson AB. Standardised classification of pre-release development in male-brooding pipefish, seahorses, and seadragons (Family Syngnathidae). BMC developmental biology. 2012;12(1):1–6.

21. Wetzel JT, Wourms JP. Embryogenesis in the dwarf seahorse, Hippocampus zosterae (Syngnathidae). Gulf and Caribbean Research. 2004;16(1):27–35.

22. Mi PT, Kornienko E, Drozdov A. Embryonic and larval development of the seahorse Hippocampus kuda. Russian Journal of Marine Biology. 1998;24(5):325–329.

23. Kornienko E. Reproduction and development in some genera of pipefish and seahorses of the family Syngnathidae. Russian Journal of Marine Biology. 2001;27(1):S15–S26.

24. Azzarello MY. A comparative study of the developmental osteology of Syngnathus scovelli and Hippocampus zosterae (Pisces, Syngnathidae) and its phylogenetic implications. Evolutionary Monographs; 1990.

25. Maters BR, Stevenson E, Vize PD. Embryonic and aglomerular kidney development in the bay pipefish, Syngnathus leptorhynchus. bioRxiv. 2021.

26. Hamilton H, Saarman N, Short G, et al. Molecular phylogeny and patterns of diversification in Syngnathid fishes. Molecular phylogenetics and Evolution. 2017;107:388–403.

27. Longo SJ, Faircloth BC, Meyer A, Westneat MW, Alfaro ME, Wainwright PC. Phylogenomic analysis of a rapid radiation of misfit fishes (Syngnathiformes) using ultraconserved elements. Molecular phylogenetics and evolution. 2017;113:33–48.

28. Braga Goncalves I, Ahnesjö I, Kvarnemo C. The relationship between female body size and egg size in pipefishes. Journal of Fish Biology. 2011;78(6):1847–1854.

29. Höch R, Schneider RF, Kickuth A, Meyer A, Woltering JM. From fin to fin - spiny and soft-rayed fin domains in acanthomorph fish are established through a BMP-gremlin-shh signaling network. PNAS. 2021.

30. Vincent AC, Berglund A, Ahnesjö I. Reproductive ecology of five pipefish species in one eelgrass meadow. Environmental Biology of Fishes. 1995;44(4):347–361.

31. Ahnesjo I. Temperature affects male and female potential reproductive rates differently in the sex-role reversed pipefish, Syngnathus typhle. Behavioral Ecology. 1995;6(2):229–233.

32. Lin Q, Fan S, Zhang Y, et al. The seahorse genome and the evolution of its specialized morphology. Nature. 2016;540(7633):395–399.

33. Monteiro N, Almada V, Vieira M. Implications of different brood pouch structures in syngnathid reproduction. Marine Biological Association of the United Kingdom Journal of the Marine Biological Association of the United Kingdom. 2005;85(5):1235.

34. Teixeira R, Musick JA. Reproduction and food habits of the lined seahorse, Hippocampus erectus (Teleostei: Syngnathidae) of Chesapeake Bay, Virginia. Revista Brasileira de Biologia. 2001;61:79–90.

35. Jones AG, Rosenqvist G, Berglund A, Avise JC. Mate quality influences multiple maternity in the sex-role-reversed pipefish Syngnathus typhle. Oikos. 2000;90(2):321–326.

36. Kawaguchi M, Okubo R, Harada A, et al. Morphology of brood pouch formation in the pot-bellied seahorse Hippocampus abdominalis. Zoological letters. 2017;3(1):19.

37. Kimmel CB, Ballard WW, Kimmel SR, Ullmann B, Schilling TF. Stages of embryonic development of the zebrafish. Developmental dynamics. 1995;203(3):253–310.

38. Iwamatsu T. Stages of normal development in the medaka Oryzias latipes. Mechanisms of development. 2004;121(7-8):605–618.

39. Parichy DM, Elizondo MR, Mills MG, Gordon TN, Engeszer RE. Normal table of postembryonic zebrafish development: staging by externally visible anatomy of the living fish. Developmental dynamics. 2009;238(12):2975–3015.

40. Walker M, Kimmel C. A two-color acid-free cartilage and bone stain for zebrafish larvae. Biotechnic & Histochemistry. 2007;82(1):23–28.

41. Leysen H, Christiaens J, De Kegel B, Boone M, Van Hoorebeke L, Adriaens D. Musculoskeletal structure of the feeding system and implications of snout elongation in Hippocampus reidi and Dunckerocampus dactyliophorus. Journal of Fish Biology. 2011;78(6):1799–1823.

42. Neutens C, Adriaens D, Christiaens J, et al. Grasping convergent evolution in syngnathids: a unique tale of tails. Journal of anatomy. 2014;224(6):710–723.

43. Lees J, Märss T, Wilson MV, Saat T, Špilev H. The sculpture and morphology of postcranial dermal armor plates and associated bones in gasterosteiforms and syngnathiforms inhabiting Estonian coastal waters. Acta Zoologica. 2012;93(4):422–435.

44. Porter MM, Adriaens D, Hatton RL, Meyers MA, McKittrick J. Why the seahorse tail is square. Science. 2015;349(6243):aaa6683.

45. Hawkins MB, Henke K, Harris MP. Latent developmental potential to form limb-like skeletal structures in zebrafish. Cell. 2021;184(4):899–911. e13.

46. Consi T, Seifert P, Triantafyllou M, Edelman E. The dorsal fin engine of the seahorse (Hippocampus sp.). Journal of morphology. 2001;248(1):80–97.

47. Kanou K, Kohno H. Early life history of a seahorse, Hippocampus mohnikei, in Tokyo Bay, Japan. Ichthyological Research. 2001;48(4):361–368.

48. Hale ME. Functional morphology of ventral tail bending and prehensile abilities of the seahorse, Hippocampus kuda. Journal of morphology. 1996;227(1):51–65.

49. Ackermann MA, Kontrogianni-Konstantopoulos A. Myosin binding protein-C slow: a multifaceted family of proteins with a complex expression profile in fast and slow twitch skeletal muscles. Frontiers in physiology. 2013;4:391.

50. Roth O, Solbakken MH, Tørresen OK, et al. Evolution of male pregnancy associated with remodeling of canonical vertebrate immunity in seahorses and pipefishes. Proceedings of the National Academy of Sciences. 2020;117(17):9431–9439.

51. Skalkos ZM, Van Dyke JU, Whittington CM. Paternal nutrient provisioning during male pregnancy in the seahorse Hippocampus abdominalis. Journal of Comparative Physiology B. 2020;190(5):547–556.

52. Otero-Ferrer F, Lättekivi F, Ord J, et al. Time-critical influences of gestational diet in a seahorse model of male pregnancy. Journal of Experimental Biology. 2020;223(3):jeb210302.

53. Kawahara R, Miya M, Mabuchi K, et al. Interrelationships of the 11 gasterosteiform families (sticklebacks, pipefishes, and their relatives): a new perspective based on whole mitogenome sequences from 75 higher teleosts. Molecular phylogenetics and evolution. 2008;46(1):224–236.

54. Dawson C. Bulbonaricus Herald (Pisces: Syngnathidae), a senior synonym of Enchelyocampus Dawson and Allen, with description of Bulbonaricus brucei n. sp. from eastern Africa. Copeia. 1984:565–571.

55. Lopez-Rios J, Duchesne A, Speziale D, et al. Attenuated sensing of SHH by Ptch1 underlies evolution of bovine limbs. Nature. 2014;511(7507):46–51.

56. Welten MC, Verbeek FJ, Meijer AH, Richardson MK. Gene expression and digit homology in the chicken embryo wing. Evolution & development. 2005;7(1):18–28.

57. Laudet V. The origins and evolution of vertebrate metamorphosis. Current Biology. 2011;21(18):R726–R737.

58. Monteiro N, Almada V, Vieira M. Early life history of the pipefish Nerophis lumbriciformis (Pisces: Syngnathidae). Journal of the Marine Biological Association of the United Kingdom. 2003;83(5):1179–1182.

59. Silva K, Monteiro N, Almada V, Vieira M. Early life history of Syngnathus abaster. Journal of Fish Biology. 2006;68(1):80–86.

60. Delunardo FAC, Paulino MG, Medeiros LCC, Fernandes MN, Scherer R, Chippari-Gomes AR. Morphological and histopathological changes in seahorse (Hippocampus reidi) gills after exposure to the water-accommodated fraction of diesel oil. Marine Pollution Bulletin. 2020;150:110769.

61. Prein M, Kunzmann A. Structural organization of the gills in pipefish (Teleostei, Syngnathidae). Zoomorphology. 1987;107(3):161–168.

62. Adams DC, Otárola-Castillo E. geomorph: an R package for the collection and analysis of geometric morphometric shape data. Methods in ecology and evolution. 2013;4(4):393–399.

63. Team RC. R: A language and environment for statistical computing. 2013.

64. Kim Y-I, Lee S, Jung S-H, et al. Establishment of a bone-specific col10a1: GFP transgenic zebrafish. Molecules and cells. 2013;36(2):145–150.

65. Ha K, Buchan JG, Alvarado DM, et al. MYBPC1 mutations impair skeletal muscle function in zebrafish models of arthrogryposis. Human molecular genetics. 2013;22(24):4967–4977.

66. Ochi H, Westerfield M. Lbx2 regulates formation of myofibrils. BMC developmental biology. 2009;9(1):1–14.

